# A Chemical Kinetic Basis for Measuring Translation Initiation and Elongation Rates from Ribosome Profiling data

**DOI:** 10.1101/490730

**Authors:** Ajeet K. Sharma, Pietro Sormanni, Nabeel Ahmed, Prajwal Ciryam, Ulrike A. Friedrich, Guenter Kramer, Edward P. O’Brien

**Author notes:** These authors contributed equally to this work (Ajeet K. Sharma, Pietro Sormanni, Nabeel Ahmed).

## Abstract

Analysis methods based on simulations and optimization have been previously developed to estimate relative translation rates from next-generation sequencing data. Translation involves molecules and chemical reactions; hence, bioinformatics methods consistent with the laws of chemistry and physics are more likely to produce accurate results. Here, we derive simple equations based on chemical kinetic principles to measure the translation-initiation rate, transcriptome-wide elongation rate, and individual codon translation rates from ribosome profiling experiments. Our methods reproduce the known rates from ribosome profiles generated from detailed simulations of translation. Applying our methods to data from *S. cerevisiae* and mouse embryonic stem cells we find that the extracted rates reproduce expected correlations with various molecular properties. A codon can exhibit up to 26-fold variability in its translation rate depending upon its context within a transcript. This broad distribution means that the average translation rate of a codon is not representative of the rate at which most instances of that codon are translated. We also find that mouse embryonic stem cells have a global translation speed of 5.2 AA/s, is similar to what has been previous reported using another analysis method. This large variability in translation rates suggests that translational regulation might be used by cells to a greater degree than previously thought.

## Introduction

Translation-associated rates influence *in vivo* protein abundance, structure and function. It is therefore crucial to be able to accurately measure these rates. The synthesis of a protein consists of three sequential phases – initiation, elongation, and termination [1–3]. During the initiation phase ribosome subunits form a stable translation-initiation complex at the start codon of the mRNA transcript [4,5]. The ribosome then stochastically moves forward along the transcript one codon at a time during the elongation phase, adding one residue to the nascent chain at each step. The elongation phase is terminated when the ribosome’s A-site reaches the stop codon, resulting in release of the fully synthesized protein. The initiation and elongation phases of translation contribute to protein levels inside a cell; indeed, alteration of their rates can cause protein abundance to vary by up to five orders of magnitude [6–8], and alter protein structure and function [9]. Termination does not influence the cellular concentration of proteins as it is not a rate limiting step [10]. Therefore, knowledge of translation initiation and codon translation rates are important to understand the regulation of gene expression.

Significant efforts have been made to extract these rates from data generated from ribosome profiling experiments [11–14], a technique that measures the relative ribosome density across transcripts [15]. In a number of methods, the actual rates are not measured but instead a ratio of rates, or other relevant quantities have been reported [16–20]. Estimates of translation-initiation and codon translation rates have helped identify the molecular determinants of these rates. For example, estimated initiation rates correlate with the stability of mRNA structure near the start codon and in the 5′ untranslated region [10,14,16,21] indicating mRNA structure can influence initiation. Similarly, codon translation rates have been found to positively correlate with their cognate tRNA abundance [16], and anti-correlate with the presence of downstream mRNA secondary structure and positively charged nascent-chain residues inside the ribosome exit tunnel [20,22]. Some of these findings are controversial as different analysis methods and data have led to contradictory results concerning the role of tRNA concentration [16,19,23,24], positively charged residues [17,20] and coding sequence (CDS) length [12,14]. Moreover, the accuracy of these methods is unknown because orthogonal, high-throughput experimental measurements of translation rates do not exist.

In the absence of data that could differentiate the accuracy of different methods, we argue that the methods most likely to be accurate will be those that are constrained by and account for the chemistry and physics of the translating system. Here, we present three such methods, derived from chemical kinetic principles that permit the extraction of translation-initiation rates, transcriptome-wide average elongation rate and individual codon translation rates from Next-Generation Sequencing (NGS) data. These methods are verified against artificial ribosome profiling data generated from detailed simulations of the translation process where the translation rates are known *a priori*. We then apply these methods to *in vivo* ribosome profiling data and extract the transcriptome-wide translation-initiation and codon translation rates in *S. cerevisiae* and transcriptome-wide average elongation rate in mouse stem cells. We show that the translation rate parameters correlate with factors known to modulate these rates, and assign absolute numbers to these rates.

## Theory

### Measuring translation-initiation rates

To derive an analytical expression relating the translation-initiation rate to the experimental observable of ribosome density along a transcript we assume steady state conditions, meaning that the ribosome flux at each codon position is equal to the rate at which fully synthesized protein molecules are released. Thus,

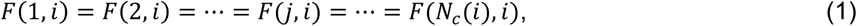

where *F*(1, *i*) is the flux of ribosomes initiating translation at the start codon, *F*(*j, i*) is the flux of ribosomes moving from codon position *j* to (*j* + 1) in transcript *i* and *N*_*c*_(*i*) is the number of codons in the transcript. In our modeling, the position of a ribosome on a transcript is defined by its A-site location.

Let ℓ be the number of codon positions that a ribosome covers on a transcript. A ribosome translating any of the first ℓ + 1 codons will prevent a new ribosome from initiating translation by physically blocking the first six codon positions of the transcript. Thus, the ribosome flux at the start codon is

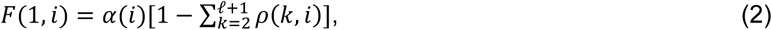

where *α*(*i*) is the initiation rate in the absence of any ribosome at the first ℓ + 1 codon positions, and *ρ*(*k, i*) is the average ribosome occupancy at the *k*^*th*^ codon position of the transcript. Since translation is initiated with the A-site of the ribosome at the second codon position, the summation in Eq. (2) starts from *k* = 2. The ribosome flux at the *j*^*th*^ codon position of transcript *i* is

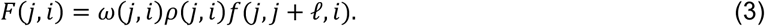

In Eq. (3), *ω*(*j, i*) is the intrinsic translation rate of the *j*^*th*^ codon position in the transcript (*i.e.*, in the absence of any other ribosomes), and *f*(*j, j* + ℓ, *i*) is the conditional probability that given that a ribosome is at the *j*^*th*^ codon position there is no ribosome at the (*j* + ℓ)^*th*^ codon position. At low ribosome densities *f*(*j, j* + ℓ, *i*) is approximately equal to (1 - *ρ*(*j* + ℓ, *i*)), which is the probability that a ribosome is not present at the (*j* + ℓ)^*th*^ codon position irrespective of the ribosome occupancy at the *j*^*th*^ codon position [25]. This assumption allows an expression for the initiation rate to be derived, but could yield inaccurate values of *F*(*j, i*)s at higher *ρ*(*j, i*)s. Polysome profiling experiments [26] and computer simulations of protein synthesis [21], however, suggest that most *S. cerevisiae* transcripts are translated in the regime of low average ribosome density. Additionally, in the simulations we describe below, the average error in the estimated *F*(*j, i*)’s (Eq. (3)) gets larger as the initiation rate increases, hitting 17.8% at an initiation rate of 0.20 *s*^−1^. 86% of *S. cerevisiae* transcripts have *in vivo* initiation rates below 0.20 *s*^−1^ [10], indicating that for most genes approximating *f*(*j, j* + ℓ, *i*) as (1 - *ρ*(*j* + ℓ, *i*)) is reasonable. Finally, the ribosome flux at the last codon position is *F*(*N*_*c*_(*i*), *i*) = *ω*(*N*_*c*_(*i*), *i*)*ρ*(*N*_*c*_(*i*), *i*).

We used Eqs. (1), (2) and (3) to derive the following expression for translation-initiation rate

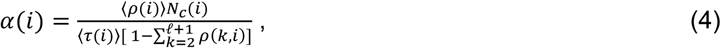

where 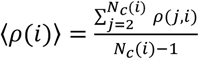 is the average ribosome density per codon on the *i*^*th*^ transcript, and 〈τ(*i*)〉 is the average time a ribosome takes to synthesize a full-length protein from the transcript. A derivation of Eq. (4) is provided in the Supporting Information (SI). Eq. (4) can measure the translation-initiation rate provided 〈τ(*i*)〉, 〈*ρ*(*i*)〉 and the *ρ*(*j, i*)s are known. We calculated 〈*ρ*(*i*)〉 and *ρ*(*j, i*) from ribosome profiling, RNA-Seq and polysome profiling data, and 〈τ(*i*)〉 using a scaling relationship [27] between protein synthesis time and CDS length. Full details for determining these parameters are provided in the SI.

### Measuring the average translation elongation rate across a transcriptome

In ribosome run-off experiments translation-initiation is inhibited at time point *t*=0 using the drug harringtonine [23]. This causes ribosomes to accumulate at start codons and thereby block new translation-initiation events. At time *t* = Δ*t*, the cells are treated with cycloheximide, which stops translation elongation by the ribosomes. The experiment is repeated at different Δ*t* values. A meta-gene analysis of ribosome profiles [28] is then obtained from each run-off experiment and used to quantify the depletion of ribosome density across the transcripts as a function of time, allowing the transcriptome-wide average elongation rate to be measured [23].

To model this non-steady state data, we assume ribosome movement along transcripts is a form of mass flow along one dimension – a reasonable assumption given that ribosomes can move only in one direction along a transcript. In this case, a natural way to describe this phenomenon is the continuity equation

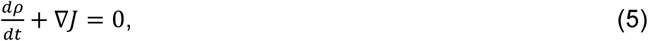

which equates the decrease in density of a substance (*ρ*) with the outward flux *J* of that substance from a particular region of space. The equality with zero in Eq. (5) is a manifestation of conservation of mass. In the ribosome run-off experiment this corresponds to a depletion of ribosome density along a segment of a transcript as ribosomes move out of that region with rate *J*, and are not replenished by new ribosomes since initiation was halted at time *t* = 0.

Solving Eq. (5) by integrating over *t* from 0 to Δ*t* yields

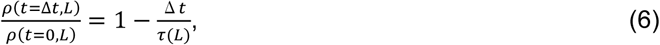

where *ρ*(*t* = Δ*t, L*) is the average ribosome density of the transcriptome within the first *L* codon positions of a sample at run-off time Δ*t*. 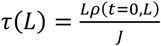 is the average time at which 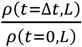 equals zero.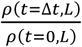 in Eq. (6) is calculated using the ribosome run-off data. Explicit details of this calculation are provided in the SI.

We inserted the relative, average ribosome density *ρ*(*t* = Δ*t, L*)/*ρ*(*t* = 0, *L*) obtained from Eq. (S10) into Eq. (6) and plotted this against Δ*t*. By fitting this data to a straight line, we determined τ(*L*) as the time at which *ρ*(*t* = Δ*t, L*)/*ρ*(*t* = 0, *L*) equals zero. We calculated τ(*L*) in this way for different *L* values up to the minimum *L* value at which ribosome density depletion no longer occurs even at the longest run off time. τ(*L*) is, therefore, the time at which the last translating ribosomes have crossed the *L*^*th*^ codon position, on average. The average transcriptome-wide elongation rate 〈*ω*〉 is therefore equal to

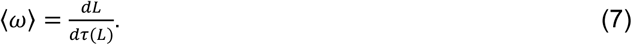

### Measuring individual codon translation rates

To derive a mathematical expression for extracting codon translation rates from ribosome profiling data we assumed steady-state conditions in which the flux of ribosomes at each codon position is equal to the rate of protein synthesis

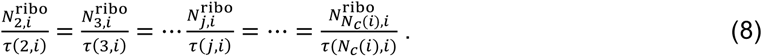

In Eq. (8), 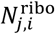 and τ(*j, i*) are the steady-state number of ribosomes and the average translation time of the *j*^*th*^ codon position within transcript *i*. The mean time of synthesis 〈τ(*i*)〉 of transcript *i* is, by definition, equal to

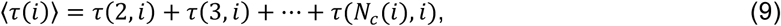

Solving Eqs. (8) and (9) for τ(*j, i*) yields

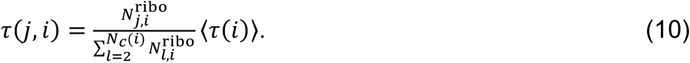

As is conventional in the field [29], we assume that ribosome profiling reads at the *j*^*th*^ codon position of transcript *i, c*(*j, i*), are directly proportional to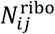. This relationship can be expressed as

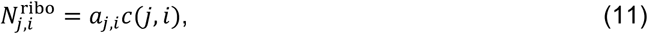

where *a*_*j,i*_ is a constant of proportionality. *a*_*j,i*_ values have not been experimentally measured, but they are commonly assumed to be constant for all codon positions in a single experiment [29]. That is, *a*_*j,i*_ = *a* for all *i* and *j*. Using Eq. (11) in Eq. (10) yields

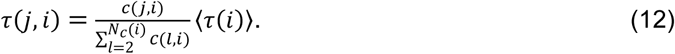

Eq. (12) indicates that we can determine the individual codon translation rates from ribosome profiling reads provided we know the average synthesis time of the transcript. Eq. (12) can be connected to the expression for normalized ribosome density, derived in the SI of Ref. [16], where 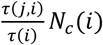 is the normalized ribosome density and is expressed as a function of *c*(*j, i*)s. Eq. (12) is also related to a metric used in the simulations of Ref. [14] to estimate the codon translation rates. It is important to note that τ(*j, i*) is the actual codon translation time which includes the time delay caused by ribosome-ribosome interactions and is therefore different than the inverse of *ω*(*j, i*).

## Computational methods

### Simulated steady state and non-steady state ribosome profiling data

1,388 *S. cerevisiae* mRNA transcripts were selected to test our translation-initiation rate measurement method. They were chosen based on the filtering criteria that at least 95% of their codons have non-zero read density in the ribosome profiling data reported in Ref. [16]. The list of these genes are provided in Supporting File 1. We used the translation-initiation rates reported in Ref. [10] in our simulations for 1,236 of the transcripts. The initiation rates for the other 152 transcripts were not reported in Ref. [10]. Therefore, we randomly assigned (with replacement) the initiation rate to those 152 transcripts from the same database. We used codon translation rates from Fluitt and Viljoen’s model for all 61 sense codons [30] and set the translation-termination rate to 35 *s*^−1^ [10]. We set ℓ = 10 codons in our simulations because it is the canonical m RNA fragment length that is protected by ribosomes against nucleotide digestion in ribosome profiling experiments [15].

We simulated the translation of these 1,388 *S. cerevisiae* mRNA sequences using the Gillespie’s algorithm [31] to generate the *in silico* ribosome profiling data. During the simulations, we recorded ribosome locations across the transcript every 100 steps, which we found minimized the time-correlation between successive saved snapshots. The codon positions of the ribosome’s A-site in each of these snapshots, summed over all snapshots, constituted the *in silico* generated ribosome profile for the transcript. We ran the simulations until the total number of *in silico* ribosome profiling reads were equal to the total number of reads aligned to the same transcript measured from experimental ribosome profiling data reported in Ref [16]. This allowed us to create a realistic level of statistical sampling in our *in silico* ribosome profiles. Each of the uncorrelated snapshots can be thought as a separate copy of the mRNA transcript. Thus, the total number of these snapshots were equal to the mRNA copy number in our *in silico* experiment which we used to calculate *ρ*(*j, i*)s.

Non-steady-state ribosome run-off experiments were simulated for 4,617 *S. cerevisiae* transcripts by using a Monte Carlo simulation method whose procedure is described in Refs. [32–34]. These transcripts were chosen based on the filtering criteria reported in Ref. [23]. To perform the *in silico* run-off experiment described in Ref. [23], first we waited until the translation system achieved a steady-state and then set *α* = 0, which stopped translation-initiation in our simulations. However, we allowed ribosomes to continue elongation for a time Δ*t*, after which elongation was halted. Next, we recorded the ribosome A-site positions across the transcript which were defined as *in silico* ribosome profiling reads. We repeated this *in silico* experiment for the run-off times of Δ*t* = 5, 10, 15, 20, 25 and 30 seconds. Note well, we did not use Gillespie’s algorithm for these simulations because each step in that algorithm occurs in a random time interval [31], preventing translation-elongation to stop at *t* = Δ*t* in our *in silico* run-off experiment.

To test our method for measuring codon translation rates we selected the transcripts which have at least three ribosome profiling reads at each codon position in the ribosome profiling data reported in Ref. [16]. 85 *S. cerevisiae* transcripts meet this criterion and are a subset of the 1,388 transcripts from our original selection. An explanation for choosing this criterion is provided later in the computational methods.

### *In silico* measurement of average protein synthesis and codon translation times

To calculate the translation-initiation rate (Eq. (4)) and codon translation times (Eq. (12)) from *in silico* ribosome profiles we need the average time a ribosome takes to synthesize a protein from a given transcript. We measured this quantity from our simulations by recording the time it takes a ribosome to traverse from the start codon to the stop codon in the transcript. The average synthesis time of a protein was then calculated from 10,000 individual ribosomes.

We also calculated the average synthesis time of a protein using a scaling relationship that uses the transcriptome-wide average codon translation time (Eq. (S8)). To calculate this quantity we first computed the average codon translation time for each transcript by dividing the average protein synthesis time of a transcript by its CDS length. We then calculated the transcriptome-wide average codon translation time using the average codon translation time of each transcript.

Testing the accuracy of Eq. (12) requires the real codon translation times which we measured by setting a separate clock at each codon position of a transcript in our simulations. These clocks measured the time difference between successive arrival and departure of a ribosome at each codon position. To calculate the average codon translation time at each codon position at least 10,000 such measurements were made.

### Calculation of the folding energy near the 5′ mRNA cap and estimation of other relevant parameters

We calculated the folding energy near the 5′ mRNA cap of *S. cerevisiae* transcripts to measure its correlation with *in vivo* translation-initiation rates calculated using Eq. (4). The folding energy of the first 70 nucleotides after the 5′ mRNA cap correlates the most with the translation-initiation rate [16]. To identify the 5′ UTR sequences included in those 70 nucleotides, we used the database reported in Ref [35]. We calculated the folding energy of each segment by using the software RNAfold 2.0 [36].

We used the mRNA and protein copy numbers reported in Ref. [37] to measure their correlation with translation-initiation rates. The mRNA and protein copy number reported in this paper are the average copy numbers, which were averaged using three independent studies [38–43].

### Analysis of ribosome profiling and RNA-Seq data

#### Datasets

To calculate the translation-initiation rates we applied our methods (Eq. (4)) to *in vivo* ribosome profiling data published in Refs. [16] and [44] with NCBI accession numbers GSM1289257 and GSM1949551, respectively. The RNA-Seq data we used for Ref. [16] has NCBI accession number GSM1969535. For Ref. [44], we make public the RNA-Seq sample at GSM3242263 which we performed simultaneously with the ribosome profiling experiment. We chose these two data sets because they contain both ribosome profiling and RNA-Seq data of a sample, allowing us to calculate the ribosome density of a transcript (Eq. (S6)). To calculate the codon translation rate, we apply our method to high-coverage ribosome profiling datasets of wild-type *S. cerevisiae* reported in Refs. [44] and [45] with NCBI accession numbers GSM1949551 and GSM1495503, respectively. In our analysis, reads were preprocessed and mapped to sacCer3 reference genome as described in Ref. [44]. To maintain the accuracy of read assignment, transcripts in which multiple mapped reads constitute more than 0.1% of total reads mapping to the CDS region were not considered in the analysis. A-site positions in ribosome profiling reads were assigned according to the offset table generated using a Linear Programming algorithm which maximizes the reads between the second and stop codon of transcripts [46]. The offset table for *S. cerevisiae* is given in Table S1. RPKM values were calculated for transcripts in RNA-Seq data by counting the reads whose 5′ ends were within the coding region of the transcript.

#### Selection of genes for translation-initiation rates

To apply our method (Eq. (4)) for extracting translation initiation rates, we restrict our analysis to genes in which 95% of codon positions of a transcript have non zero read density. This threshold reduces the statistical uncertainty in the estimation of codon position dependent ribosome density used in Eq. (4).

#### Selection of genes for codon translation rates

To extract individual codon translation rates, we restrict our analysis to genes that have at least 3 reads at every codon position of the transcript. We find that 118 and 364 genes meet this criterion in the data set of Nissley and co-workers [44] and Weissman and co-workers [47], respectively. This stringent requirement is necessary since Eq. (12) would predict codons with zero reads to be translated in zero time. Reads at the start codon and the second codon have contributions from the translation initiation process; therefore we ignored these codon positions in our calculations. As stated before, transcripts containing multiple mapped reads greater than 0.1% of the total reads mapped to the transcript were discarded. Genes with overlap of coding sequence regions as well as those containing introns (which is less than 6% of *S. cerevisiae* genome) were not considered in the analysis to avoid overlap of ribosome profiles.

#### Preparation of RNA-Seq sample

200 mL of cells were grown in YPD to an OD_600 nm_ of 0.5, filtered (All-Glass Filter 90mm, Millipore), flash frozen in liquid nitrogen and lysed by mixer milling (2 min, 30 Hz, MM400, Retsch) with 600 µL of lysis buffer (20 mM Tris-HCl pH 8.0, 140 mM KCl, 6 mM MgCl2, 0.1% NP-40, 0.1 mg/ml CHX, 1 mM PMSF, 2x protease inhibitors (Complete EDTA-free, Roche), 0.02 U/ml DNaseI (recombinant DNaseI, Roche), 20 mg/mL leupeptin, 20 mg/mL aprotinin, 10 mg/mL E-64, 40 mg/mL bestatin). Thawed cell lysates are cleared by centrifugation (20,000xg, 5 min, 4°C) and RNA extraction is performed as described in Ref. [48]. 10 µg of extracted RNA was depleted for rRNA using the kit RiboMinus, Yeast Module (Invitrogen) and fragmented for 30 min using the NEBNext Magnesium RNA Fragmentation Module (NEB). Deep sequencing libraries were prepared following the protocol described in Ref. [48] and sequenced on a HiSeq 2000 (Illumina).

#### Miscellaneous

(a) Since experimental measurements of τ(*i*)s are not available for *S. cerevisiae* we use Eq. (S8) to estimate τ(*i*) with 〈τ^*A*^〉 = 200 *ms* [49,50]. (b) *In vivo* ribosome profiling data for the mouse stem cells were processed using the method described in the original paper [23]. (c) The measures for tRNA abundance based on gene copy number and RNA-Seq measurements were obtained from Table S2 of Ref. [16]. (d) Transcript leader (or 5′ UTR) sequences were obtained from Ref. [35] and upstream AUG were identified in these sequences for the transcripts for which we calculate the translation initiation rates. (e) For Kozak sequence analysis, 12 nt sequence was identified around the start codon starting from −6 with respect to Adenine of start codon (position +1) to +6. The consensus Kozak sequence for *S. cerevisiae* is (A/T)A(A/C)A(A/C)AATGTC(T/C) [51]. For the 12 nt sequence for every transcript, a similarity score is calculated based on its match with the Kozak sequence. The score ranges from 1 to 10 in the order of its increasing similarity with Kozak sequence. If all the 9 nt around the start codon are same as the Kozak sequence and the start codon (scored as 1), the score will be 9+1=10. If only start codon is same while all other 9 nt are different, the score is just 1. If start codon is same and only 4 nt positions are same, then the score will be 4+1=5. To determine the effect of Kozak sequence, two subsets of transcripts were created, the first with transcripts having context around start codon closer to Kozak sequence (score > 7) and the second subset for transcripts with context farther away from Kozak sequence (score < 5). Mann-Whitney U test was used to determine the statistical significance of the difference between the translation initiation rate distributions of the two subsets of transcripts.

### Assignment of mRNA secondary structure

Both DMS and PARS data provide information about base-paired nucleotides within an mRNA molecule. We considered a codon to be structured if at least two of its three nucleotides were base-paired or one nucleotide was base-paired and the structure information for the other two nucleotides were not available.

DMS data for *S. cerevisiae* were downloaded from GEO database with accession number GSE45803 [52]. The reads from all replicates were pooled together and then aligned to the ribosomal RNA sequences using Bowtie 2 (v2.2.3) [53]. The reads which did not align to the ribosomal RNA sequences were then aligned to the *Saccharomyces cerevisiae* assembly R64-2-1 (UCSC: sacCer3) using Tophat (v2.0.13) [54]. In our analysis, A and C nucleotides were considered base-paired when the DMS signal was below the threshold of 0.2 and considered unstructured if the DMS signal was greater than 0.5. A and C nucleotides with DMS signal between 0.2 and 0.5 are considered ambiguous and classified together with U and G nucleotides, which do not react with DMS. Codons involving such nucleotides were not considered in our analysis.

PARS data were downloaded from genie.weizmann.ac.il/pubs/PARS10 with PARS scores available for all transcripts, except YDR461W and YNL145W, which were excluded from our analysis. Nucleotides with a PARS score greater than 0 were considered base-paired [55].

## Results

### *In silico* validation of our methods

As a first step to test the accuracy of the measured translation rates from Eqs. (4), (7) and (12), we applied them to *S. cerevisiae* ribosome profiles generated by kinetic Monte Carlo simulations in which all of the underlying rates are known (see Methods). If our analysis methods are accurate then a necessary condition is that they reproduce these rates. We applied Eq. (4) to the simulated, steady-state ribosome profiling data and find that it quantitatively reproduces the initiation rates used in the simulations (Fig. 1(A), slope=0.97, *R*^2^ =0.84 and p-value ≤ 3 × 10^−308^). We applied Eq. (7) to the non-steady state ribosome run off profiles of *S. cerevisiae* (Figs. 2(A) and 2(B)) and find that it accurately measures the transcriptome-wide average translation rate. Specifically, our method measures the transcriptome-wide average elongation rate at 3.8 AA/s against the real average elongation rate of 4.2 AA/s. Note well, that as predicted by Eq. (6), we also observe a linear decrease in 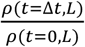 as a function of time (Fig. S1(A)). And finally we applied Eq. (12) to the steady-state ribosome profiles and find that the individual codon translation times are accurately measured by our method (Fig. 3, median *R*^2^ = 0.96 and median slope =1.00). Thus, the analysis methods we have created can in principle accurately reproduce translation rate parameters.

**Figure 1:**
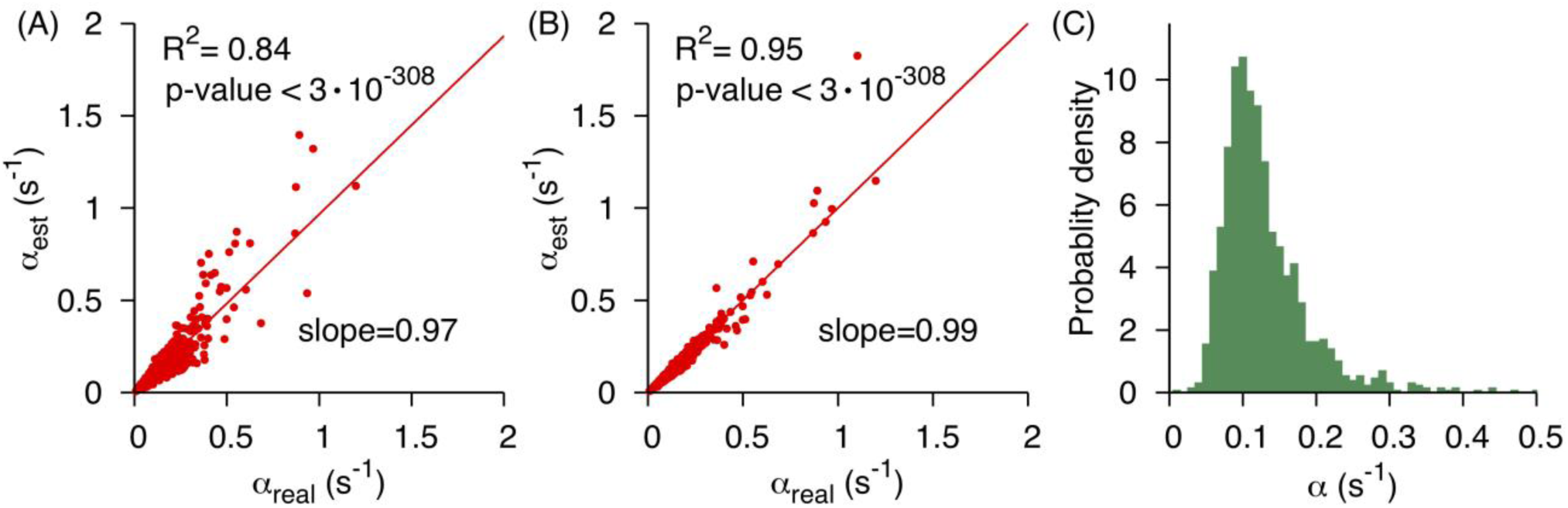
Eq. (4) accurately determines the translation-initiation rate from simulated *S. cerevisiae* ribosome profiles. (**A**) Translation initiation rates determined by applying Eq. (4) to simulated ribosome profiling data are plotted against the actual initiation rates used in the simulations. These initiation rates were calculated using Eq. (S8) for the protein synthesis times. (**B**) Same as (**A**) but the average protein synthesis times were measured from our simulations of the translation process. The solid lines in (**A**) and (**B**) are the lines of the best fit. (**C**) The distribution of *in vivo* translation-initiation rates measured by applying Eq. (4) to 1,287 *S. cerevisiae* high coverage transcripts.

**Figure 2:**
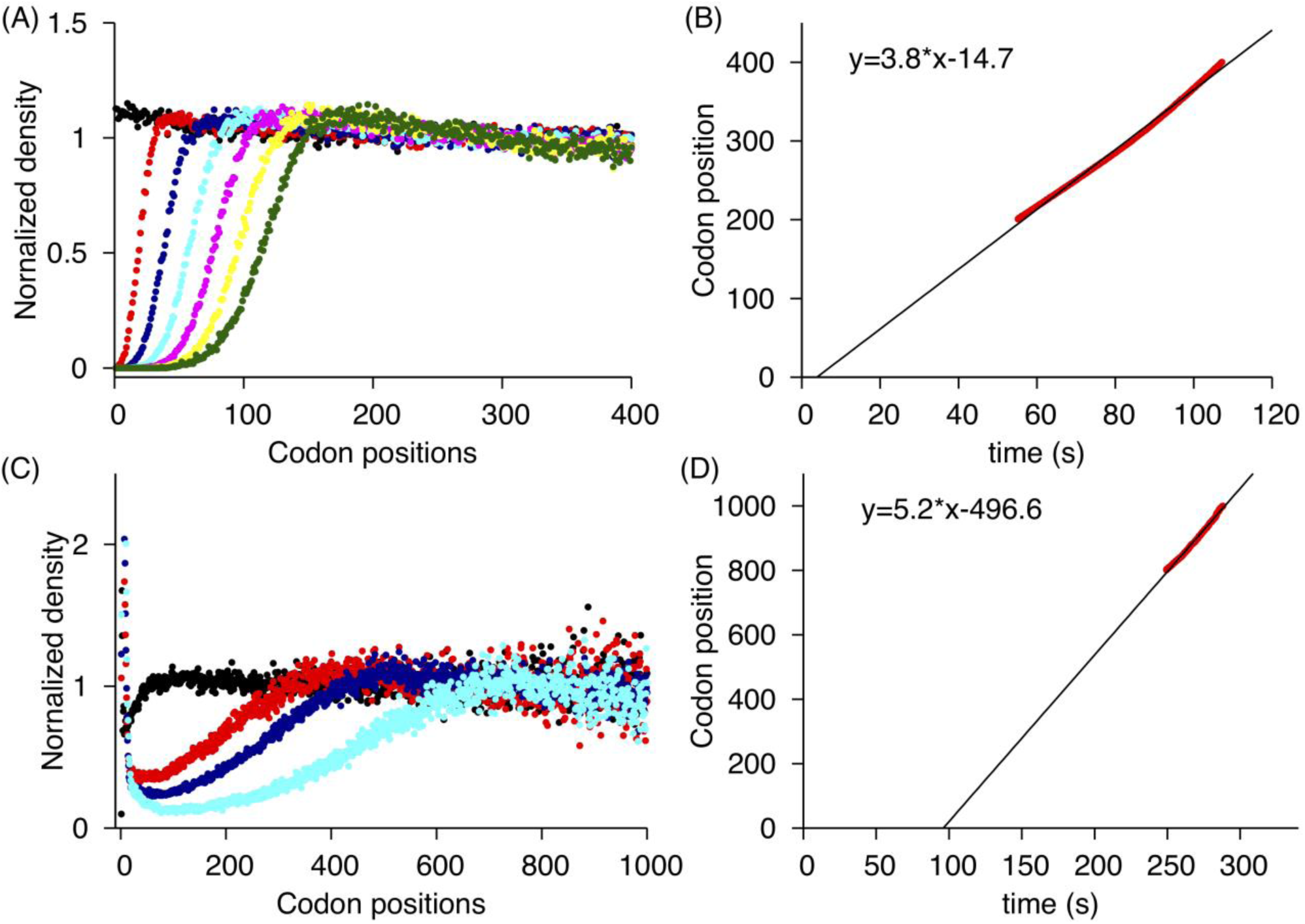
Measuring average elongation speed by applying Eq. (7) to *in silico* and *in vivo* ribosome run-off data. (**A**) Normalized average ribosome read density 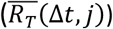, Eq. (S9), calculated from simulated ribosome run-off experiment is plotted as a function of codon position for run-off times of 0, 5, 10, 15, 20, 25 and 30 *s*^−1^ with black, red, blue, cyan, pink, yellow and green data points, respectively. (**B**) The average time taken to fully deplete the normalized average ribosome read density within a window of the most 5′ codons positions in *S. cerevisiae* transcripts are plotted against the most 3′ codon positon of the window. (**C**) Normalized average ribosome read density, calculated from *in vivo* run-off experimental data reported in Ref. [23], are plotted as a function of codon position for the run-off times of 0, 90, 120 and 150 seconds with black, red, blue and cyan data points, respectively. (**D**) The average time taken to fully deplete the normalized average ribosome read density within a window of the most 5′ codons positions in mouse stem cells transcripts are plotted against the most 3′ codon positon of the window. The negative intercept reflects the time taken by harringtonine to engage with ribosomes.

**Figure 3:**
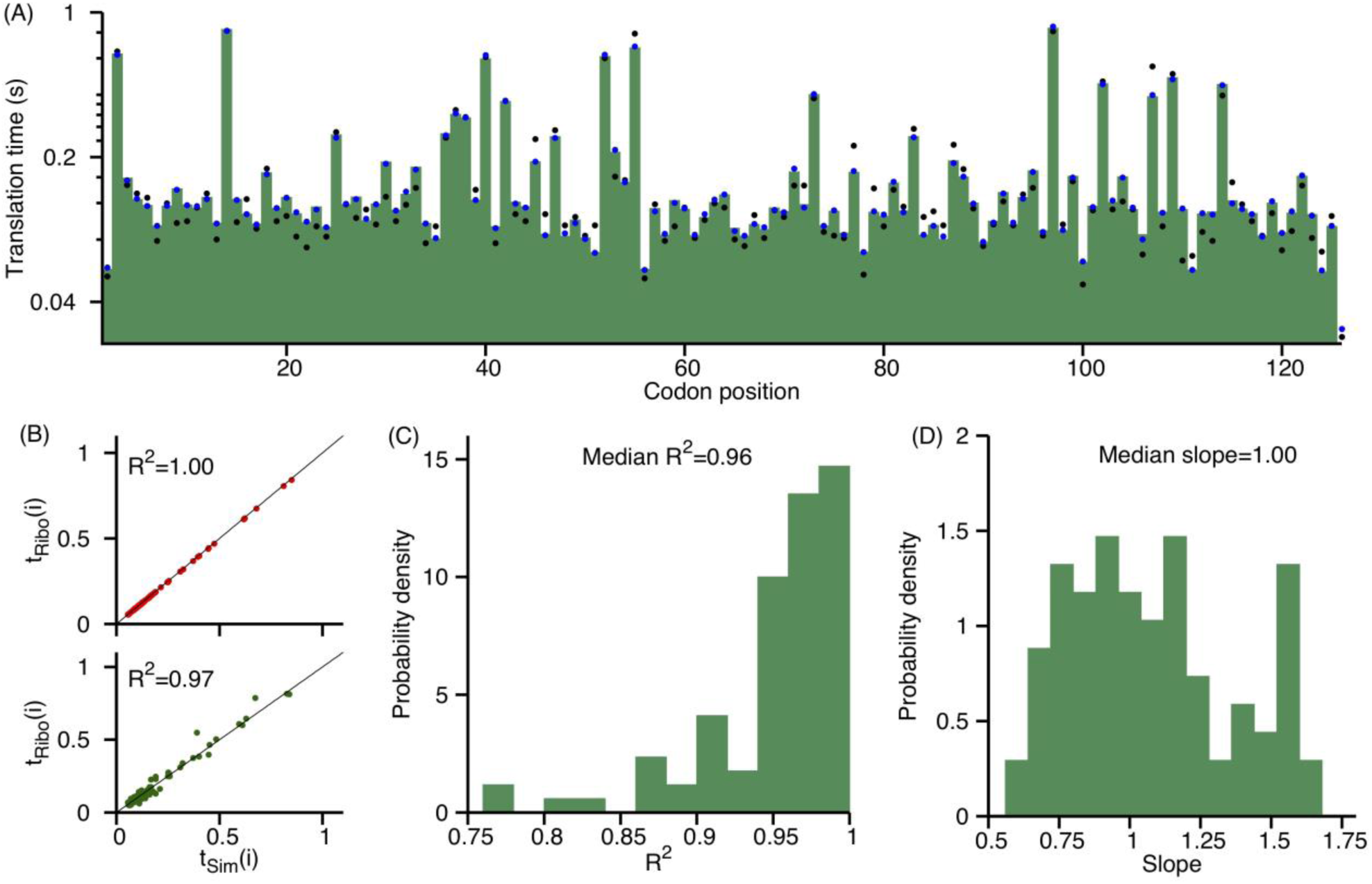
Eq. (12) accurately determines codon translation times from simulated ribosome profiles. (**A**) Average translation time of a codon in YER009W *S. cerevisiae* transcript is plotted as a function of its position within the transcript. The true codon translation times in the simulations are plotted as green boxes, blue and black data points are the translation times measured using Eq. (12). Blue data points were calculated using the average protein synthesis time measured from the simulations and relative ribosome density calculated using a large number of *in silico* ribosome profiling reads. Black data points were calculated using the average protein synthesis time estimated from the scaling relationship (Eq. (S8)) and the relative ribosome density calculated from the *in silico* reads which were equal to the reads aligned to the same transcript in the experiment [16]. (**B**) Measured codon translation times, plotted with black and blue data points in (**A**), are plotted against true codon translation times in the simulations in the top and bottom panel, respectively. (**C**) Probability distribution of the *R*^2^ correlation between the true and calculated codon translation times for the 85 *S. cerevisiae* transcripts. (**D**) Probability distribution of the slope of the best fit lines between the estimated and true codon translation times for the 85 *S. cerevisiae* transcripts. The high *R*^2^ in (**C**) and median slope of 1.00 in (**D**) indicate that Eq. (12) can, in principle, accurately measure absolute rates.

There are several points worth noting concerning these tests. First, the rates used in the simulation model are realistic, having been taken from literature values [10,30]. Second, the depth of coverage in the simulated ribosome profiles is in the same range as experiments, *e.g.*, having 26 million reads arising from 1,388 different coding sequences [16]. Third, Eqs. (4) and (12) require knowledge of the average synthesis time of a protein, which is experimentally difficult to measure. Therefore, in the above analyses we used the approximation that the average synthesis time of a protein is equal to the number of codons in coding sequence of its transcript, multiplied by the transcriptome-wide average codon translation rate (Eq. (S8)) [23,27]. However, when we use the actual synthesis time of each transcript we achieve even better agreement between the estimated and true translation initiation rates (Fig. 1B, slope = 0.99, *R*^2^ = 0.95 and p-value ≤ 3 × 10^−308^). Similarly, when we increase our read coverage from 7.1 million to 35.5 billion reads and use the exact synthesis time of a protein, *R*^2^ between the estimated and true codon translation rates goes to > 0.99 for all 85 transcripts. Thus, the methods are reasonably accurate when approximate protein synthesis times are used (Eq. (S8)) and highly accurate when the exact synthesis time is used and coverage is high.

### Measuring the initiation rate from experimental data

We next applied Eq. (4) to two different *in vivo* ribosome profiling and RNA-Seq data from *S. cerevisiae* reported in Refs. [16,44]. Calculation of the translation-initiation rate for a transcript requires knowledge of the average time a ribosome takes to synthesize the protein (〈τ(*i*)〉) and the average occupancies of the first ten codon positions (*ρ*(*j, i*)). The average protein synthesis time was calculated using the scaling relationship (Eq. (S8)) where the transcriptome-wide average codon translation time was chosen as 200 ms [49,50] (see SI). *ρ*(*j, i*)s were calculated from Eq. (S7) using the ribosome profiling and RNA-Seq data from Ref. [16] and polysome profiling data from Ref. [56]. We then inserted these arguments into Eq. (4) to calculate the translation-initiation rates for 1,287 *S. cerevisiae* transcripts which meet the filtering criteria (see Methods) for which *in vivo* average ribosome density data were available. We find the translation-initiation rates vary from as low as 5.8 × 10^−2^ *s*^−1^ (5^th^ percentile) to as high as 0.24 *s*^−1^ (95^th^ percentile) with the most probable rate being 0.1 *s*^−1^ (Fig. 1(C) and Supporting file 1). These translation-initiation rates are significantly lower than the average elongation rate in *S. cerevisiae* which is consistent with previous studies indicating that initiation is the rate limiting step in translation [21]. The distribution of translation-initiation rates measured using Eq. (4) is very similar to the distribution reported in Ref. [10], which was calculated using polysome profiling data.

### Reproducing known correlations with initiation rates

Next, we tested whether our measurements reproduce previously reported correlations between initiation rates and CDS length, sequence context upstream and around the start codon, folding energy near the 5′ cap of mRNA molecule, and protein copy number. Initiation rate may depend on CDS length because the shorter a transcript, the more probable the termini will be in close proximity, allowing more efficient re-initiation of terminating ribosomes [57]. We used the initiation rates extracted from ribosome profiling and RNA-Seq data reported in Ref. [16] and polysome profiling data reported in Ref. [56] and find a moderate but statistically significant correlation (Pearson *r* = 0.51 p-value = 4 × 10^−59^) between the translation-initiation rate and the inverse of CDS length (Fig. 4(A)), as has been observed previously [10,14,16].

**Figure 4:**
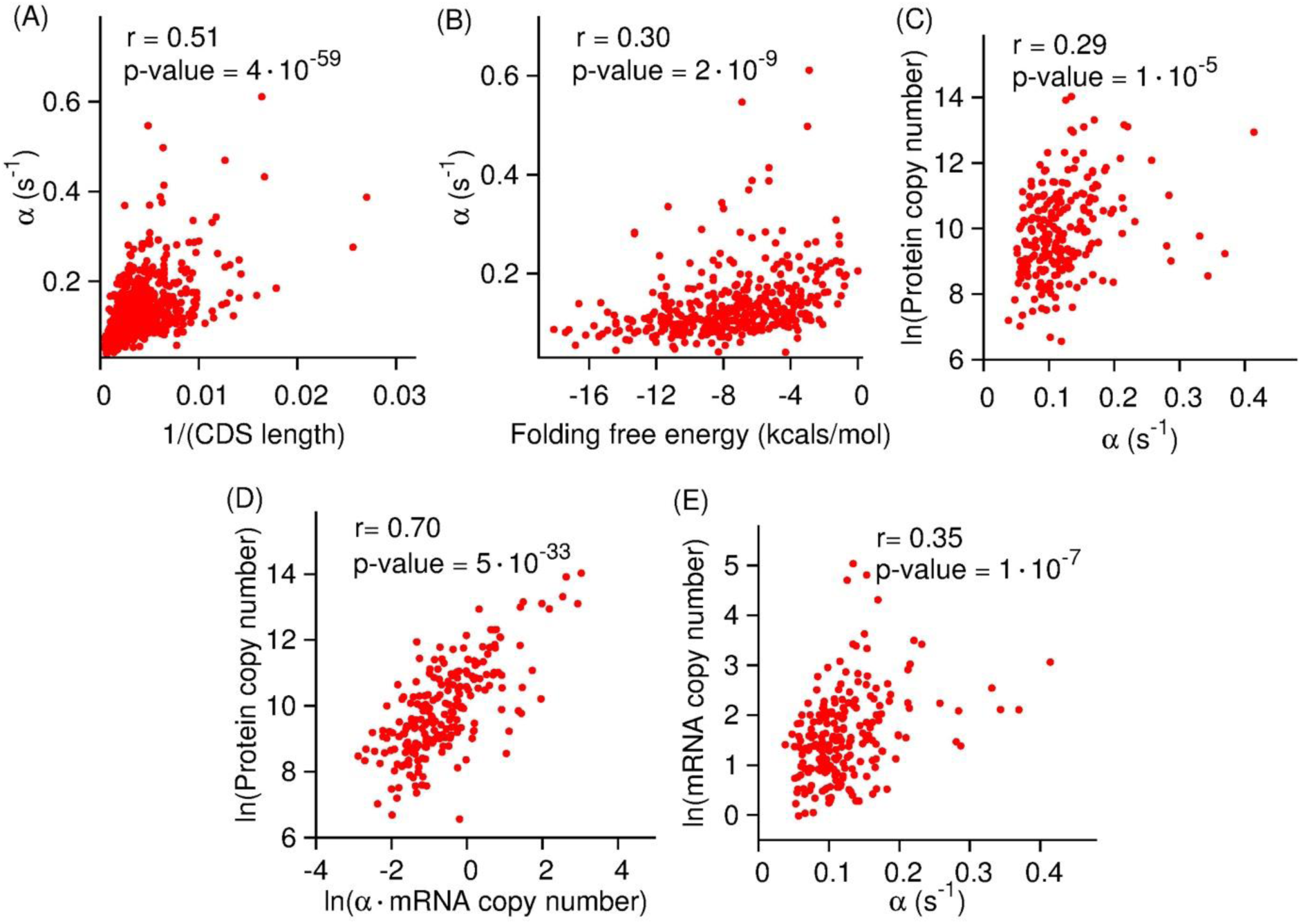
Translation-initiation rates measured using Eq. (4) reproduce previously reported correlations with molecular properties. *In vivo* translation-initiation rates of *S. cerevisiae* transcripts are plotted against the inverse of their CDS length, folding energy of mRNA molecule near the 5′ cap and protein copy number in (**A**), (**B**) and (**C**), respectively. (**D**) The copy number of *S. cerevisiae* proteins are plotted as a function of the product of the initiation rate of the transcripts that encode them and that transcript’s copy number in a cell. (**E**) mRNA copy number is plotted against the translation-initiation rate.

Folded mRNA structure near the 5′ cap can disrupt the binding of initiation factor eLF4F and the scanning of 40S ribosomes in eukaryotes, causing a decrease in a gene’s translation-initiation rate [14,16]. We calculated the mRNA folding energy near the 5′ mRNA cap (see Methods) and find a statistically significant correlation between them and our translation-initiation rates (Fig. 4(B), Pearson *r* = 0.30 and p-value = 2 × 10^−9^).

Sequence-based features also determine the rate of translation-initiation in eukaryotes. For example, an upstream open reading frame can potentially interfere with translation initiation at the canonical start codon by initiating translation prematurely resulting in a different protein product [58,59]. We tested whether upstream open reading frames affect initiation rates by comparing the median initiation rate in transcripts with at least one upstream AUG site, against transcripts that do not contain any upstream AUG. We find that the transcripts with at least one upstream AUG codon have an 15% slower median initiation rate of 0.095 *s*^−1^ as compared to the others where it is 0.112 *s*^−1^ (Mann-Whitney U test, p-value = 0.006). Weinberg and co-workers [16] also demonstrated this effect with their measure of initiation efficiency. Additionally, the 12-nt Kozak sequence [51] is enriched in highly expressed genes and mutations in the Kozak sequence have been shown to drastically effect protein abundance levels [60]. We find that initiation rates are 16% faster at 0.116 *s*^−1^ in transcripts with Kozak-like sequences around the start codon as compared to those that do not contain these mRNA sequence motifs (Mann-Whitney U test, p-value = 0.005). Hence, the presence of Kozak, or Kozak-like sequences around the start codon facilitate translation initiation. This is consistent with results of Koller and co-workers [13] who have shown a strong correlation between the presence of the Kozak sequence with of translation efficiency.

Initiation is the rate limiting step of translation [21]. Therefore, a faster initiation rate should increase the rate of protein synthesis and thus the steady-state level of proteins in a cell. Indeed, we find a statistically significant correlation between the rate of translation-initiation and protein copy number in a cell (Fig. 4(C), Pearson *r* = 0.29 and p-value = 1 × 10^−5^). Next, we tested whether the missing variation in Fig. 4(C) is explained by the variation in mRNA copy number. We did this by measuring the correlation between the protein copy number and the multiplicative product of a transcript’s copy number and its initiation rate. We find a stronger correlation between this quantity and protein copy number (Fig. 4(D), Pearson *r* = 0.70 and p-value= 5 × 10^−33^). These results suggest, as has been found previously [21,61], that the translation-initiation rate is a major determinant of protein abundance inside a cell.

We further measured the correlation between the mRNA copy number and initiation rates and found a moderate level of correlation between them (Fig. 4(E), Pearson *r* = 0.35 and p-value = 1 × 10^−7^). This suggests that higher copy number transcripts have a faster translation-initiation rate, which is consistent with previous studies suggesting that nature has two independent mechanisms to increase the copy number of highly-expressed proteins [62]. We also find statistically significant correlations between the aforementioned quantities when we calculated the translation-initiation rate using all possible combinations of ribosome profiling, RNA-Seq [16,44] and polysome profiling datasets [26,56] (Figs. S2-S4, Tables S2-S3).

In summary, we find that the initiation rates we measured correlate with factors that have been established to influence translation speed. This suggests our initiation rates are reasonable.

### Measurement of the average codon translation rate in a cell

Next, we applied Eq. (7) to measure the average codon translation speed inside mouse stem cells from ribosome run-off experiments [23]. We calculated the average normalized reads 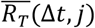 from three different samples prepared by allowing the ribosomes to continue their elongation for 90, 120, and 150 seconds after the inhibition of initiation, and plotted 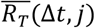 as a function of codon position (Fig. 2(C)). By following the analysis procedure described in the Methods section we fitted 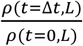 against Δ*t* curves (Fig. S1(B)) with a line and calculated the depletion time τ(*L*), where 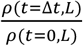 equals to zero for *L* varying between codon positions 800 and 1000. Plotting τ(*L*) against *L* yields a straight line (Fig. 2(D)). The gradient of this line is the average elongation rate, which we find equals 5.2 AA/s.

### Measurement of individual codon translation rates

To extract individual codon translation times along a coding sequence we applied Eq. (12) to 118 and 364 high-coverage transcripts from ribosome profiling data reported, respectively, in Refs. [44] and [47] (see Methods). The number of transcripts in both of these datasets are small as compared to the size of *S. cerevisiae* transcriptome. Therefore, to determine whether these subsets of transcripts are representative of the whole transcriptome we compared the distributions of different physicochemical properties in these two sets to the total transcriptome. We find that the subset of transcripts from Nissley and co-workers [44] have 6.6% higher mean GC content but a very similar mode of length distribution and codon usage, relative to the total transcriptome (Figs. S5 (A)-(C)). For Weissman and co-workers’ ribosome profiling dataset [47], again we find the mode of the length distribution and codon usage similar to the *S. cerevisiae* transcriptome with a 5.0% higher mean GC content (Figs. S5 (D)-(F)). This indicates that our transcript sets are largely representative of the properties of the transcriptome but have a bit higher GC content.

Upon extracting individual codon translation times from these ribosome profiling data, we first characterized the distribution of translation times for the 61 sense codons (Supporting file 1). We find around three-fold difference between the median translation times of the fastest and slowest codon. The fastest and slowest codons are AUU and CCG codons that are translated in 127±2 and 349±43 ms (median ± standard error), respectively. The variability in translation times for a given codon type is even larger, as illustrated by wide distributions of their translation times (Fig. 5(A)). Fig. 5(B) shows an example where AAG codon is translated with translation times ranging from 59 ms at position 413 to 363 ms at position 196 in YAL038W *S. cerevisiae* transcript. We find a 16-fold variability in codon translation times across the transcriptome even if we ignore the extremities of the distributions by only considering the translation times between the 5^th^ and 95^th^ percentiles of all codon types. Similar ranges are found in the Weissman dataset where there is a 26-fold variability in translation times and 3.9 fold-difference in median translation times of the fastest (AUC) and slowest (CGA) codons which are translated with median time of 128±2 and 497±61 ms, respectively (Supporting file 1).

**Figure 5:**
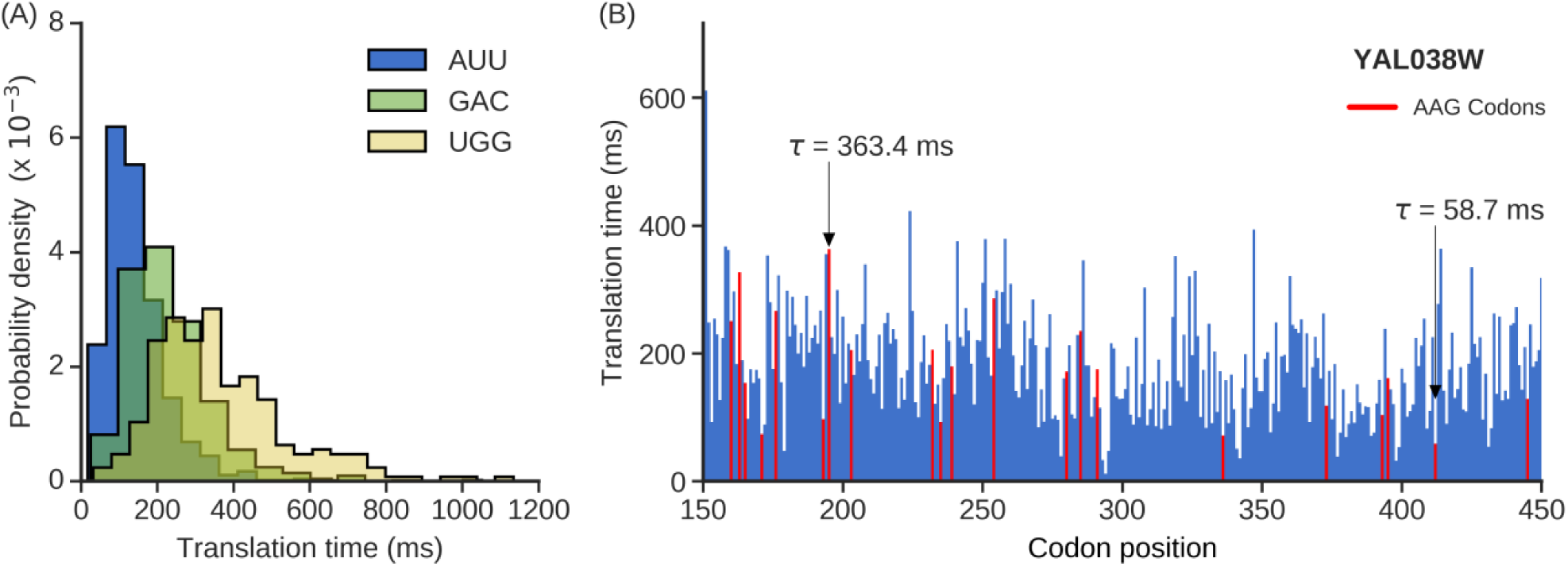
Wide variability in individual codon translation rates *in vivo*. **(A)** Probability density functions for translation times of AUU, GAC and UGG codons. Median translation times for AUU, GAC and UGG codon are 127, 209 and 334 ms, respectively. (**B**) The translation time profile *of S. cerevisiae* transcript YAL038W is shown between codon positions 150 and 450. AAG codon (colored red) is translated in 363.4 ms at the 196^*th*^ codon position and in 58.7 ms at 413^*th*^ codon position.

### Molecular factors flanking the A-site shape the variability of individual codon translation rates

A number of molecular factors have been shown or implicated to influence the translation rate of a codon in the A-site, including tRNA concentration, mRNA structure, wobble-base pairing, and proline residues at or near the ribosome P-site [16,18–20,63]. Influence of these molecular factors by varying strength is responsible for the high variation in the translation rate of a codon type at different places. Here, we test whether the presence or absence of these factors correlate with changes in translation speed. We first examined whether the cognate tRNA concentration correlates with our measured translation times. We found that the median codon translation times negatively correlates with the abundance of cognate tRNA (Figs. 6(A) and (B), *ρ* = −0.50 (p-value = 8 × 10^−4^) and *ρ* = −0.50 (p-value= 0.001), respectively). This shows that a codon is typically translated in a lesser time if its cognate tRNAs are available with higher abundance.

**Figure 6:**
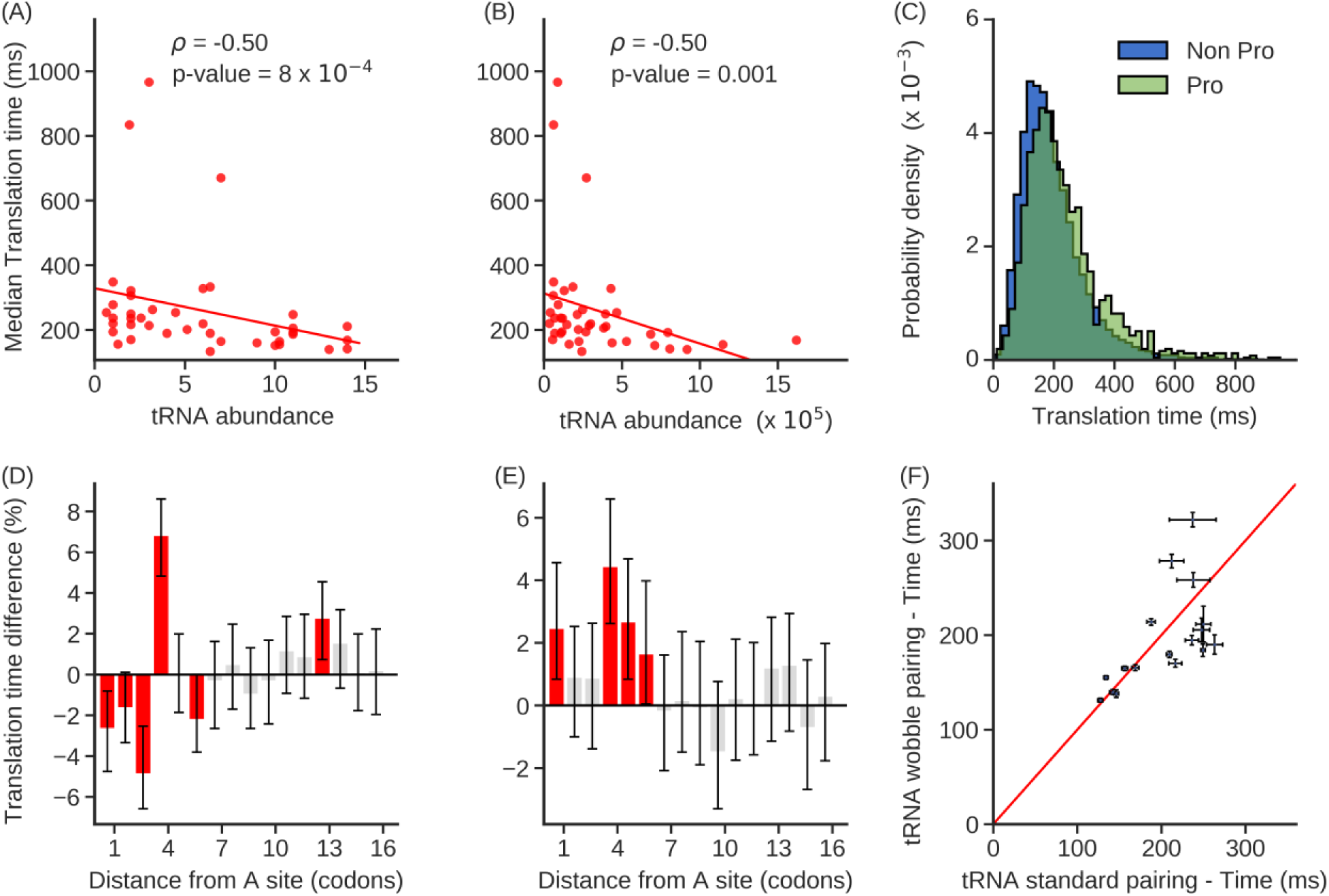
Molecular factors shaping the variability of individual codon translation rates. **(A-B)** Median translation times of codon types are negatively correlated with cognate tRNA abundance estimated by **(A)** gene copy number and **(B)** RNA-Seq gene expression. **(C)** Probability distribution of the translation time of codons which are followed by the proline encoding codon and the rest of the other codons are plotted in green and blue, respectively. **(D-E)** Percentage difference in median translation times when mRNA structure is present relative to when it is not present is plotted as a function of codon position after the A-site. Grey bars indicate results that are not statistically significant. Error bars are the 95% C.I. calculated using 10^4^ bootstrap cycles; significance is assessed using the Mann-Whitney U test corrected with the Benjamini Hochberg FDR method for multiple-hypothesis correction. mRNA structure information used in (**D**) and (**E**) are provided by *in vivo* DMS and *in vitro* PARS data, respectively. **(F)** Scatter plot of the median translation times of pairs of codon types that are decoded by the same tRNA molecule. The red line is the identity line. The list of tRNA molecule names and decoded codon types were taken from Ref. [79]. Error bars are standard error about the median translation time calculated with 10^4^ bootstrap cycles.

The presence of a proline amino acid at the ribosome’s P-site can slowdown translation due to its stereochemistry [64]. We tested whether such an effect was present in our data set by calculating the percentage difference in median translation time at the A-site when proline is present at the P-site versus when it is not. We find a 19.1% increase in median translation time when proline is present (Fig. 6(C), Mann-Whitney U test, p-value = 8.7 × 10^−33^).

It has been found that the presence of downstream mRNA secondary structure can slow down the translation at A-site [20,65]. To test for this effect, we plotted the difference in the median translation time at the A-site when mRNA secondary structure is present and when it is not present at a given number of codon positions downstream to the ribosome A-site. Structured versus unstructured nucleotides were identified using DMS data [52]. We find that when secondary structure is present 4 codons downstream of the A-site, placing that structure directly at the leading edge of the ribosome, there is on average a 6.8% increase in codon translation time at the A-site (Fig. 6(D), Mann-Whitney U test, p-value = 2.7 × 10^−14^). A similar result is also found when we cross-reference our codon translation times with mRNA structure data from PARS [55], which measures the presence of mRNA structure *in vitro* (Fig. 6(E), Mann-Whitney U test, p-value = 5.6 × 10^−9^).

Wobble base pairing between the codon and anti-codon tRNA stem-loop has been found to slowdown translation speed as compared to Watson-Crick base pairing in bacteria [66] and metazoans [67]. For each pair of codon types that are decoded by the same tRNA molecule, by Watson-Crick base pairing in one instance and wobble base pairing in the other, we tested whether two codon types are translated with different rates. We find that there is no systematic difference in median translation times between codons that are decoded by either mechanism (Fig. 6(F), Wilcoxon signed-rank test, p-value=0.46), indicating that, at least in *S. cerevisiae*, wobble base pairing does not slowdown *in vivo* translation elongation.

These results were reproduced using another dataset [47] which also shows that codon translation times anti-correlate with tRNA concentration (Figs. S6(A) and S6(B), *ρ* = −0.58 (p-value = 7.8 × 10^−5^) and *ρ* = −0.56 (p-value = 0.0002), respectively), exhibit significant slow-down in codon translation time when a proline is present at the P-site (Fig. S6(C), Mann-Whitney U test, p-value = 3.0 × 10^−27^) and mRNA structure present downstream to the A-site (Figs. S6(D) and S6(E) Mann-Whitney U test, p-values = 3.6 × 10^−5^, 8.7 × 10^−4^, respectively) and similarly, we found no difference between the translation rate of codons that are translated with Watson-Crick and Wobble base pairing (Fig. S6(F), Wilcoxon signed-rank test, p-value=0.88).

## Discussion

We have presented three methods for measuring initiation rates and elongation rates from ribosome profiling data. What distinguishes our approach from many others is that it uses simple equations derived from chemical kinetic principles, it does not require simulations or a large number of parameters, and it yields absolute rather than relative rates. We demonstrated that our approach provides accurate results when applied to test data sets (Figs. 1, 2 and 3), and reproduced previously reported correlations between translation speed and various molecular factors (Figs. 4, 6, S2-S4 and S6).

Two novel findings concerning elongation rates are that in *S. cerevisiae* the translation time of a codon depends dramatically on its context within a transcript; and that the average translation rate of a codon type is not representative of the rate at which most instances of that codon are translated. In *S. cerevisiae*, the range of individual codon translation time spans up to 26-fold, from 45 to 1197 ms, even after ignoring the extremities of the distribution of codon translation times. The codon AAG in gene YAL038W for example occurs 36-times along the transcript. At the 196^th^ codon position it is translated in 363 ms, and at the 413^th^ position it is translated in 59 ms. Thus, the same codon in different contexts can be translated at very different speeds. Characterizing the distribution of mean times of translation of different occurrences of the same codon reveals a broad distribution (Fig. 5(A)), whose coefficient of variation is often close to 0.5 (Supporting file 1). This means that the standard deviation is half of the average of this distribution. Therefore, the average translation time of a codon type is not representative of most of the times in which that codon is actually translated. These results are consistent with the findings that a large number of molecular factors determine codon translation rates [68], giving rise to a broad distribution (Fig. 5(A)). Eq. (7) was shown to accurately determine the average elongation rate from simulated data sets (Fig. 2(B)). Applying Eq. (7) to ribosome run-off experiments revealed that in mouse stem cells the average codon translation rate is 5.2 AA/s that is similar to the elongation rate measured in a previous study [23].

Measuring initiation rates using Eq. (4) reproduced previously reported correlations (Figs. 4(A)-(C)), and also revealed a statistically significant correlation between the rate of translation-initiation and mRNA copy number (Fig. 4(E)). This correlation indicates that genes with a higher mRNA copy number tend to have a higher translation efficiency, suggesting that transcriptional and translational regulation of gene expression can act synergistically to maximize the protein copy number of highly expressed genes.

Other methods have determined initiation rates by varying the initiation rate in simulations until the average ribosome density of a transcript matched the experimentally measured value [10,12,14,69]. These methods require knowledge of individual codon translation rates. Thus, small errors in these rates have the potential to accumulate and lead to large errors in the estimated initiation rates. This might contribute to conflicting conclusions reported in the literature. For example, Song and co-workers [14] found a negative correlation of initiation rate with CDS length, whereas de Ridder and co-workers [12] found no such correlation. In contrast to these methods, our method is based on a simple chemical kinetic equation that is easy to implement and does not require any detailed information of codon translation rates. Apart from these simulation-based approaches, other methods [19,21,70,71] measure the protein synthesis rate, which represents a lower bound to the rate of translation-initiation (Eq. (2)). One of these methods uses chemical kinetic modeling [21], but unlike Eq. (4), this method requires a number of biophysical parameters, including the diffusion constant of tRNAs and ribosomes, and cell volume. The difference between the initiation rate and protein synthesis rate increases with increasing initiation rate [72]. Therefore, such methods could exhibit greater errors for transcripts that have higher translation-initiation rates, which are often found in highly expressed transcripts (Figs. 4(C) and 4(E)).

A number of approaches have been developed to measure codon translation times including simulation based approaches [12,14] that extract rates by comparing the local distribution of ribosome profiling reads with simulated ribosome densities, others that optimize an objective function [13] or fit a normalized-footprint-count distribution of a codon to an empirical function [11], and yet others that measure relative codon translation times by quantifying the enrichment of ribosome read density using a variety of procedures [18,19]. In contrast, Eq. (12) allows individual, absolute codon translation rates to be calculated directly from the ribosome profile along the transcript. Another distinction is that a number of these methods [11–13,19] assume that all occurrences of a codon across the transcriptome must be translated at the same rate. This assumption cannot be correct as it is known that non-local aspects of translation (such as mRNA structure) can influence the translation speed of individual codons. Eq. (12) does not make this assumption, and therefore its extracted rates will better reflect the naturally occurring variation of codon translation times across a transcript.

Ribosome profiles have ill-quantified sequencing biases that can potentially produce reads that are not proportional to the underlying number of ribosomes at a particular codon position. This could lead to errors in the extraction of translation rate parameters using our methods [73]. Experimental improvements that minimize bias have been developed [29,63,74,75], such as utilizing short microRNA library generation techniques [76], but sequence-dependent biases can still exist, for example due to varying efficiencies of linker ligation [77]. With knowledge of the percent excess or depletion of reads over the true ribosome density we could add and subtract reads to a ribosome profile at each codon position and correct for such sequencing biases. However, these experimental measurements and data do not exist to our knowledge. While sequencing bias appears to be negligible in our data – as we see clear biological signals from the determinants of translation speed - when such data becomes available the accuracy of our extracted rates can be further improved.

The absence of accurate translation rate parameters is an impediment to quantitatively modeling the translation process. By measuring translation rate parameters using a chemical-kinetic framework, our methods have the potential to contribute to ongoing efforts [71,78] to understand how the sequence features of an mRNA molecule can regulate gene expression. More broadly, the approach we have taken in this study is to utilize ideas from chemistry and physics to analyze Next-generation Sequencing data; a branch of bioinformatics we refer to as physical bioinformatics. We expect that this physical-science-based approach will prove useful in understanding other large biological data sets, and compliment the conventional computer science approaches to bioinformatics.

## Supporting information

## Author Contributions

A.K.S, P.S., P.C. and E.P.O. conceived the study. N.A., A.K.S, P.S., P.C. and E.P.O. contributed to design of computational analyses. U.A.F and G.K. performed the RNA-Seq experiment and provided the experimental data. A.K.S, P.S., P.C. and N.A. analyzed the data. A.K.S, N.A., P.S., P.C. and E.P.O wrote the manuscript. All authors reviewed and commented on the manuscript.

## Acknowledgments

P.S. is also supported by a Borysiewicz Biomedical Fellowship from the University of Cambridge. E.P.O. acknowledges support from an NSF CAREER Award and NIH MIRA R35.

