## Supplementary material for "A Chemical Kinetic Basis for Measuring Translation Initiation and Elongation Rates from Ribosome Profiling data"

**Supplementary Methods**

**Derivation for the analytical expression for the translation-initiation rate.** We derived an expression for translation-initiation rate under steady-state conditions (Eq. (1)). To do this, we equated Eqs. (2) and (3), which is valid according to Eq. (1), and then solved for $\omega(j,i)$, yielding

$$\omega(j,i) = \frac{\alpha(i)\left[1-\sum_{k=2}^{\ell+1}\rho(k,i)\right]}{\rho(j,i)[1-\rho(j+\ell,i)]}\,. \tag{S1}$$

Excluded volume interactions between ribosomes translating the same mRNA molecule increases the average amount of time they spend at each codon position. Therefore, the effective rate at which the $j^{th}$ codon position is translated by a ribosome decreases from $\omega(j,i)$ to $\omega(j,i)[1-\rho(j+\ell,i)]$ because a ribosome sitting at the $(j+\ell)^{th}$ codon position blocks the forward movement of the ribosome at the $j^{th}$ codon position. Thus, the average time, $\langle\tau(i)\rangle$, a ribosome takes to translate transcript $i$, composed of $N_c(i)$ codons, is

$$\langle\tau(i)\rangle = \sum_{j=2}^{N_C(i)} \frac{1}{\omega(j,i)[1-\rho(j+10,i)]}\,, \tag{S2}$$

where $\rho(j+\ell,i) = 0$ when $j > N_C(i) - 10$. Substituting $\omega(j,i)$ from Eq. (S1) into Eq. (S2), and solving for $\alpha(i)$ yields

$$\alpha(i) = \frac{<\rho(i)>N_C(i)}{<\tau(i)>[\,1-\sum_{k=2}^{\ell+1}\rho(k,i)]}\,, \tag{4}$$

where $\langle\rho(i)\rangle = \frac{\sum_{j=2}^{N_C(i)}\rho(j,i)}{N_C(i)-1}$ is the average ribosome density of the $i^{th}$ transcript.

**Estimation of $\langle\boldsymbol{\rho(i)}\rangle$, $\boldsymbol{\rho(j,i)s}$ and $\langle\boldsymbol{\tau(i)}\rangle$.** Measuring translation-initiation rates using Eq. (4) requires knowledge of $\langle\rho(i)\rangle$, $\rho(j,i)s$ and $\langle\tau(i)\rangle$, which we calculated using a combination of ribosome profiling, RNA-Seq and polysome profiling data. To calculate $\langle\rho(i)\rangle$ we used the experimental observation that the number of RNA-Seq reads aligned to a transcript is proportional to that transcript's copy number ($n_m(i)$) and coding sequence length [1]. We also make the conventional assumption that the number of ribosome profiling reads aligned to a transcript is proportional to the total number of ribosomes ($n_R(i)$) translating that transcript [2]. Therefore,

$$d(i) = a_1 n_m(i) N_c(i) \tag{S3}$$

and

$$c(i) = a_2 n_R(i), \tag{S4}$$

where $d(i)$ and $c(i)$ are the reads aligned to transcript $i$ in RNA-Seq and ribosome profiling experiments, respectively, and $a_1$ and $a_2$ are proportionality constants. Dividing Eq. (S4) by (S3) yields the average ribosome density per codon on transcript $i$

$$\langle\rho(i)\rangle = \frac{n_R(i)}{n_m(i)N_c(i)} = \frac{a_1}{a_2}\frac{c(i)}{d(i)}\,. \tag{S5}$$

The ratio $\frac{c(i)}{d(i)}$ in Eq. (S5) is proportional to translation efficiency ($TE(i)$), defined as the ratio of ribosome profiling to RNA-Seq experiment reads in units of per kilobase, per million reads mapped to transcript $i$ [3]. Thus,

$$\langle\rho(i)\rangle = \xi\, TE(i), \tag{S6}$$

where $\xi$ is a function of $a_1$, $a_2$ and total number of reads aligned in the ribosome profiling and RNA-Seq experiment.

As pointed out in Ref. [4], the $\xi$ can be determined from the best fit line to the $\langle\rho(i)\rangle$ from polysome profiling (Ref. [5]) versus the $TE(i)$ calculated from ribosome profiling and RNA-Seq data (Ref. [3]). We carried out this analysis and find a statistically significant correlation between $\langle\rho(i)\rangle$ and $TE(i)$ (Fig. S7(A), Pearson $r = 0.51$, p value $\leq 3\times10^{-308}$) with $\xi = 0.015$. Similar values of $\xi$ are found using all combinations of ribosome profiling and RNA-Seq data [6] as well as polysome profiling data [7] (Fig. S7). With this value of $\xi$, we can use Eq. (S6) to calculate $\langle\rho(i)\rangle$ for any transcript, even those not in the original polysome profiling data set.

The $\rho(j,i)$s can be calculated by multiplying the probability that given a ribosome is translating transcript $i$ it will be found at codon positon $j$ (*i.e.*, $\frac{c(j,i)}{c(i)}$) by the average number of transcript $i$ (*i.e.*, $\langle\rho(i)\rangle N_c(i)$)

$$\rho(j,i) = \frac{c(j,i)}{c(i)}\langle\rho(i)\rangle N_c(i). \tag{S7}$$

We estimated the synthesis time of a protein by using the finding that it scales linearly with the number of codons in a transcript [8]

$$\langle\tau(i)\rangle \;=\; N_c(i)\langle\tau^{\mathrm{A}}\rangle. \tag{S8}$$

In Eq. (S8), $\langle\tau^{\mathrm{A}}\rangle$ is the transcriptome-wide average codon translation time. This approximation is supported both by experimental results [9] and a theoretical analysis that indicates this estimate is typically within 5% of the true synthesis time [8].

Thus, all the terms on the right hand side of Eq. (4) can be determined by utilizing data from ribosome profiling, RNA-Seq and polysome profiling, allowing for the determination of the initiation rate of each transcript.

**Estimation of $\frac{\rho(t=\Delta t,L)}{\rho(t=0,L)}$ for use in Eq. (6).** Calculation of the transcriptome-wide average elongation rate requires the use of relative ribosome density $\frac{\rho(t=\Delta t,L)}{\rho(t=0,L)}$ in Eq. (6), which we calculated from the meta-gene analysis of the ribosome run-off experimental data. To do that first we calculated the average ribosome density at each codon position as

$$R_T(j,\Delta t) = \frac{\sum_i R(i,j,\Delta t)}{N_t(j)}\,, \tag{S9}$$

where $R(i,j,\Delta t)$ is the ribosome profiling reads aligned at codon position $j$ on transcript $i$ divided by the average ribosome profiling reads per codon in that transcript in a sample with the run-off time $\Delta t$, and $N_t(j)$ is the number of mRNA transcripts identified in the experiment whose CDS length is either equal or greater than $j$ codons. The total number of aligned ribosome profiling reads in a sample can vary from one experiment to the next, therefore we further normalized $R_T(j,\Delta t)$ by dividing this to its average value calculated in the region of the meta-gene profile in which no depletion of ribosome reads occurs at the longest run-off time. For example, we normalized $R_T(j,\Delta t)$ in mouse embryonic stem cells runoff data by dividing to the average value of $R_T(j,\Delta t)$ between codon positions 800 and 1,000 (Fig. 2).

We assume that the number of ribosome profiling reads aligned to a transcript is proportional to the number of ribosomes translating that transcript. Under this assumption, the number of normalized average ribosome profiling reads $\overline{R_T}(j,\Delta t)$ within the first $L$ codon positions of the transcripts is proportional to $\rho(t=\Delta t,L)$. Therefore,

$$\frac{\rho(t=\Delta t,L)}{\rho(t=0,L)} = \frac{\sum_{j=2}^{L}\overline{R_T}(j,\Delta t)}{\sum_{j=2}^{L}\overline{R_T}(j,0)}. \tag{S10}$$

This expression for $\frac{\rho(t=\Delta t,L)}{\rho(t=0,L)}$ is then used in Eq. (6) to calculate the transcriptome-wide average elongation rate.

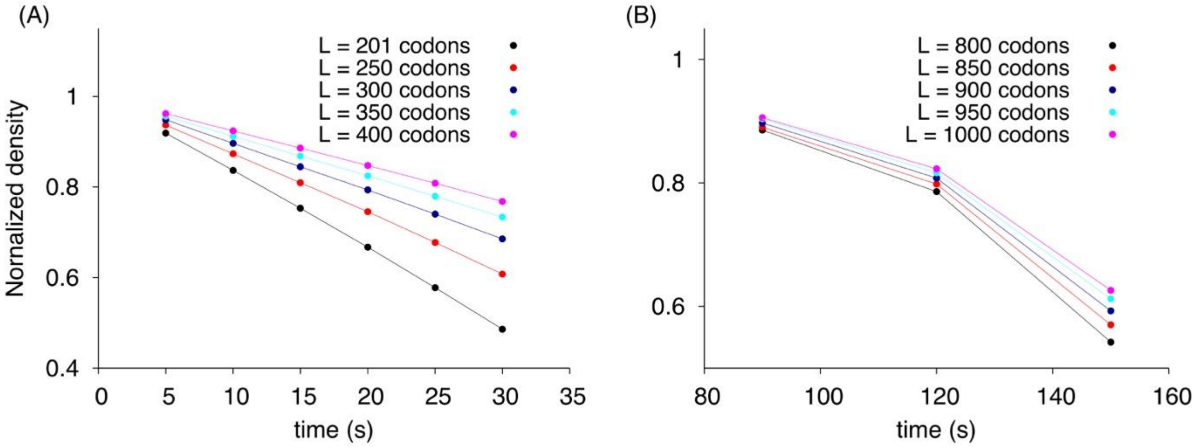


**Figure S1**: **Normalized ribosome read density calculated from run-off experiments decreases linearly as a function of time**. (**A**) The normalized ribosome read density in *S. cerevisiae* using *in silico* run-off experiment data in the first 201, 250, 300, 350 and 400 codons are plotted as a function of time. (**B**) The normalized ribosome read density in mouse stem cells (Ref. [9]) are plotted as a function of time in the first 800, 850, 900, 950 and 1000 codons. Lines are to guide the eye.

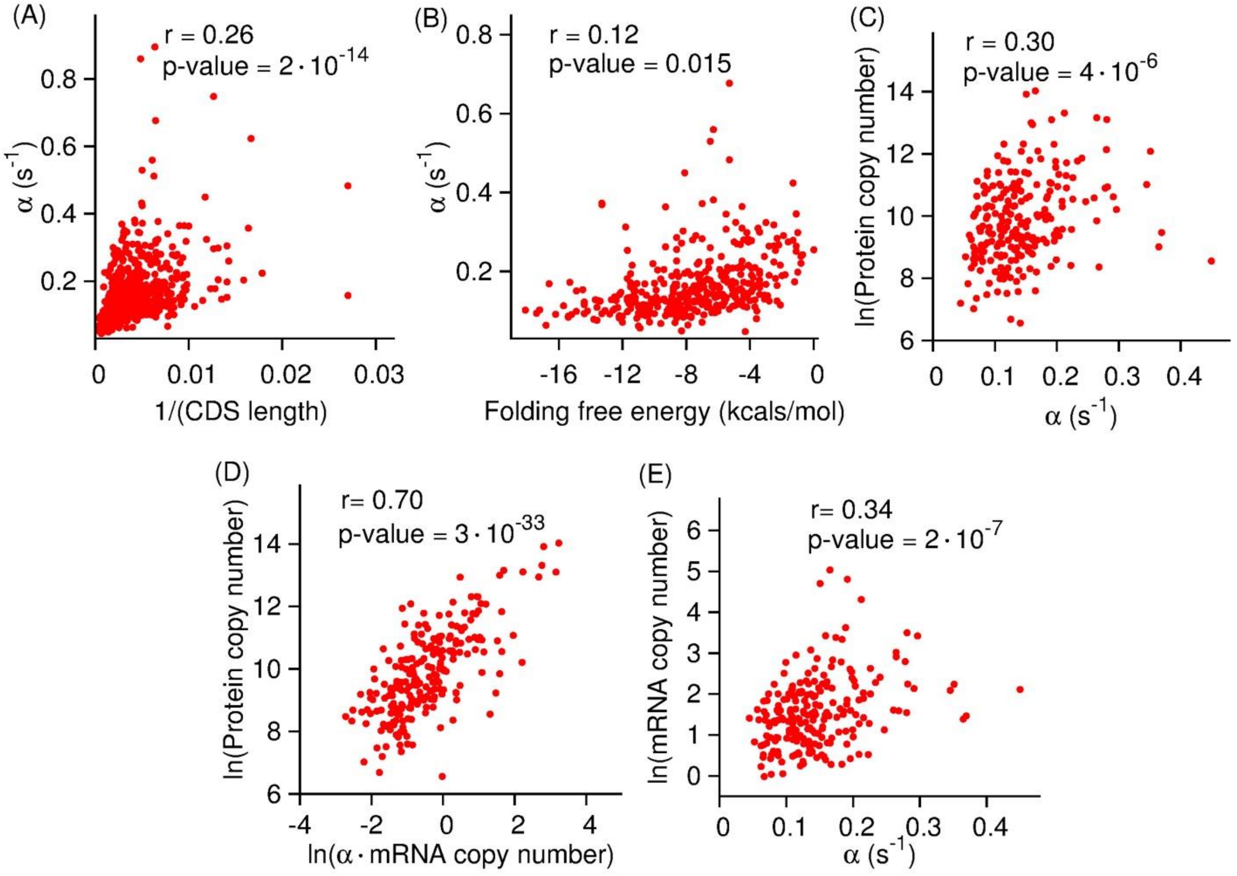


**Figure S2**: **Translation-initiation rates measured from data sets taken from Refs.** [3] **and** [7] **reproduce previously reported correlations.** *In vivo* translation-initiation rates of *S. cerevisiae* transcripts are plotted against the inverse of their CDS length, folding energy of mRNA molecule near the 5′ cap and protein copy number in (**A**), (**B**) and (**C**), respectively. (**D**) The copy number of *S. cerevisiae* proteins are plotted as a function of the product of the initiation rate of transcripts that encode them and that transcript's copy number in a cell. (**E**) mRNA copy number is plotted against the translation-initiation rate.

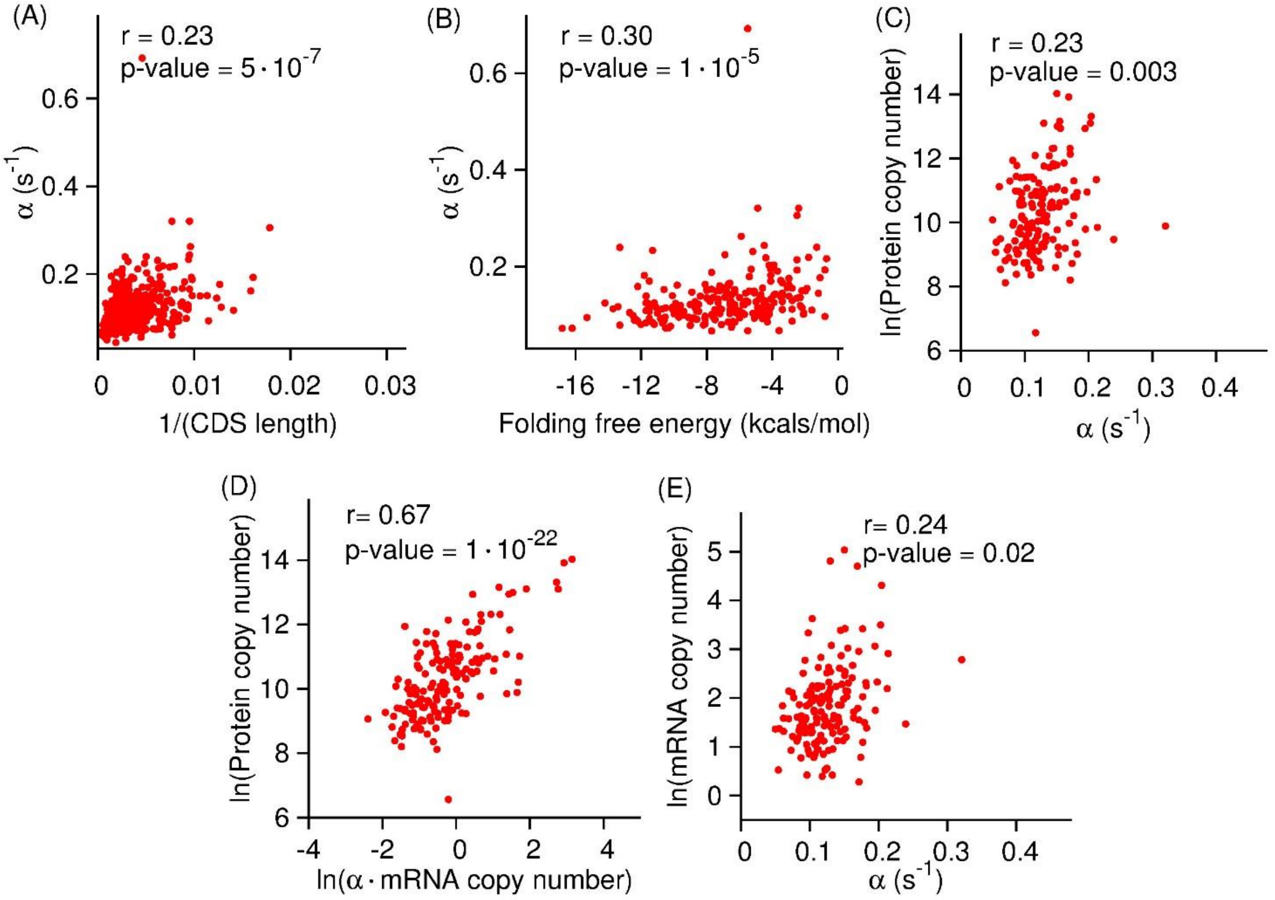


**Figure S3**: **Translation-initiation rates measured from data sets taken from Refs.** [6] **and** [5] **reproduce previously reported correlation.** *In vivo* translation-initiation rates of *S. cerevisiae* transcripts are plotted against the inverse of their CDS length, folding energy of mRNA molecule near the 5′ cap and protein copy number in (**A**), (**B**) and (**C**), respectively. (**D**) The copy number of *S. cerevisiae* proteins are plotted as a function of the product of the initiation rate of the transcripts that encode them and that transcript's copy number in a cell. (**E**) mRNA copy number is plotted against the translation-initiation rate.

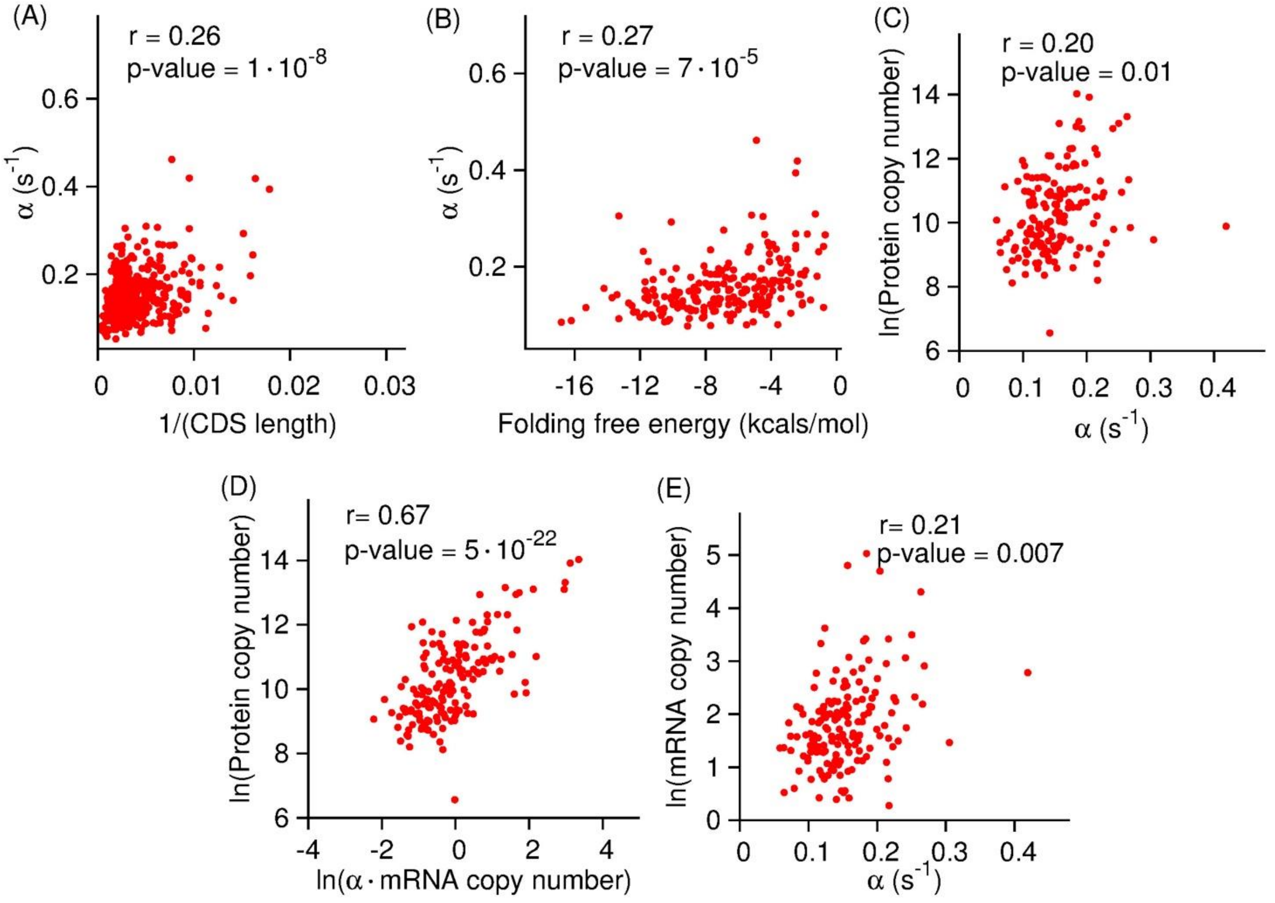


**Figure S4**: **Translation-initiation rates calculated from data sets taken from Refs.** [6] **and** [7] **reproduce previously reported correlations.** *In vivo* translation-initiation rates of *S. cerevisiae* transcripts are plotted against the inverse of their CDS length, folding energy of mRNA molecule near the 5′ cap and protein copy number in (**A**), (**B**) and (**C**), respectively. (**D**) The copy number of *S. cerevisiae* proteins are plotted as a function of the product of the initiation rate of the transcripts that encode them and that transcript's copy number in a cell. (**E**) mRNA copy number is plotted against the translation-initiation rate.

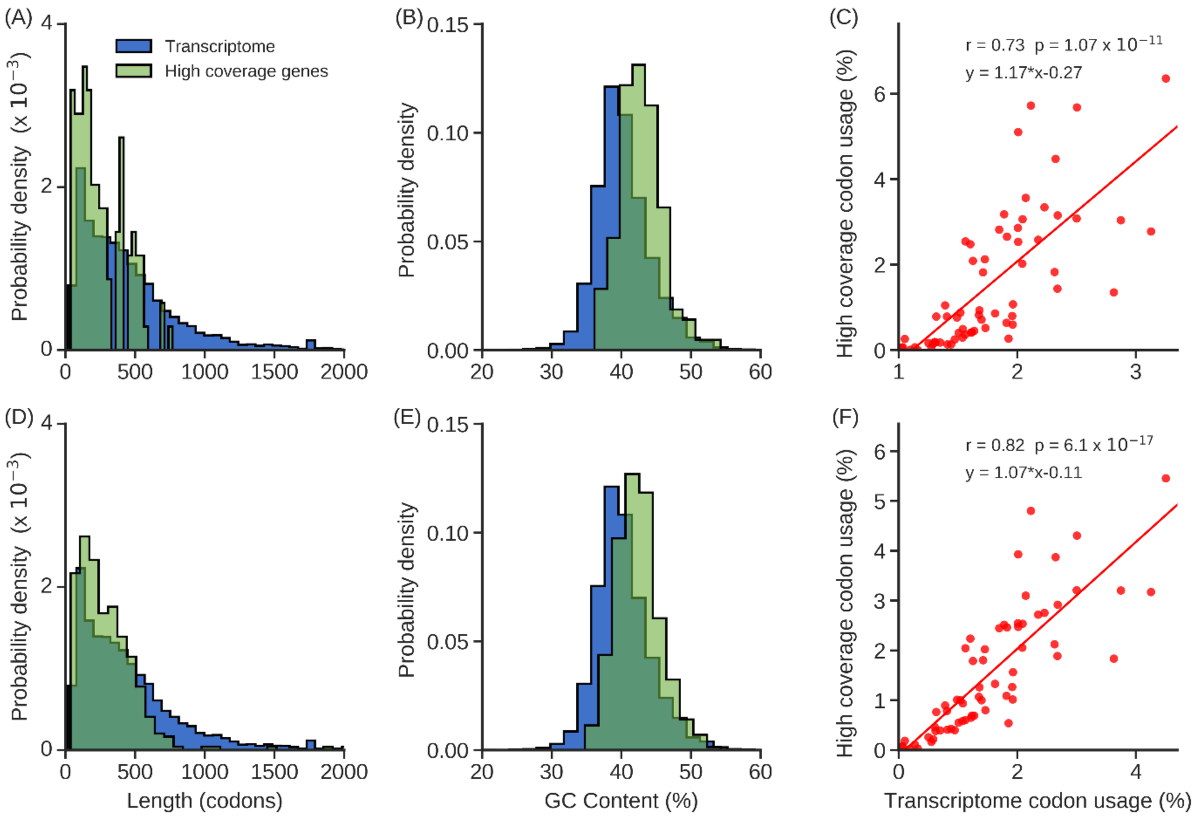


**Figure S5: Comparison of the properties of the 118- and 364-transcript data sets from Refs.** [6] **and** [10]**, respectively, to the entire *S. cerevisiae* transcriptome.** Probability distributions of CDS length and percent GC content from the data set of 118-transcripts from Ref. [6] (green) and from the entire transcriptome (blue) are plotted in (**A**) and (**B**), respectively. (**C**) Scatter plot of the codon usage in the whole genome versus the 118-transcript data set from Ref. [6]. (**D**), (**E**) and (**F**) are the same as (**A**), (**B**) and (**C**), respectively, except 364-transcripts from Ref. [10] is used**.**

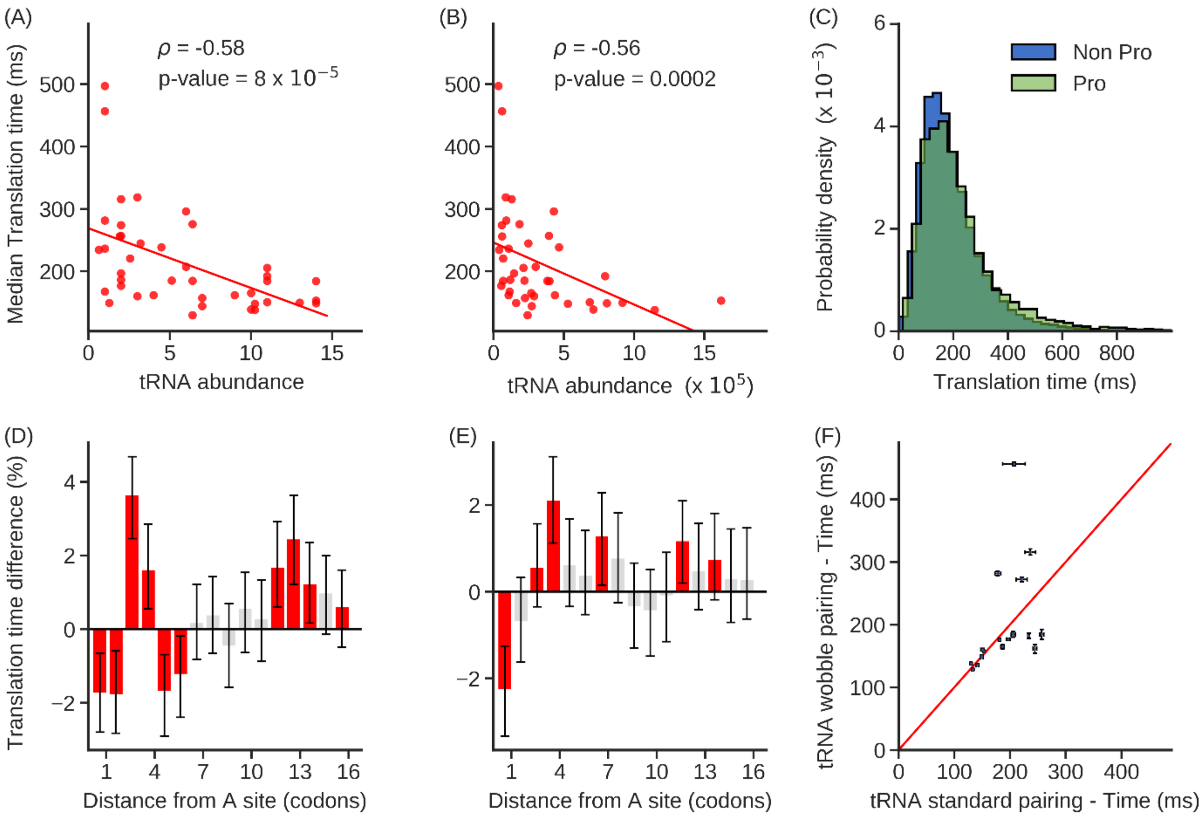


**Figure S6: Molecular factors shaping the variability of individual codon translation rates in the dataset from Ref** [10]**. (A-B)** Median translation times of codon types are negatively correlated with cognate tRNA abundance estimated by **(A)** gene copy number and **(B)** RNA-Seq gene expression. **(C)** Probability distribution of the translation time of codons which are followed by the proline encoding codon and the rest of the other codons are plotted in green and blue, respectively. **(D-E)** Percentage difference in median translation times when mRNA structure is present relative to when it is not present as a function of codon position after the A-site. Grey bars indicate results that are not statistically significant. Error bars are the 95% C.I. calculated using $10^4$ bootstrap cycles; significance is assessed using the Mann-Whitney U test corrected with the Benjamini Hochberg FDR method for multiple-hypothesis correction. mRNA structure information used in (**D**) and (**E**) were taken from *in vivo* DMS and *in vitro* PARS data, respectively. **(F)** Scatter plot of the median translation times of pairs of codon types that are decoded by the same tRNA molecule. The red line is the identity line. The list of tRNA molecules and which codon they decode were taken from Ref. [11]. Error bars are standard error about the median calculated with $10^4$ bootstrap cycles.

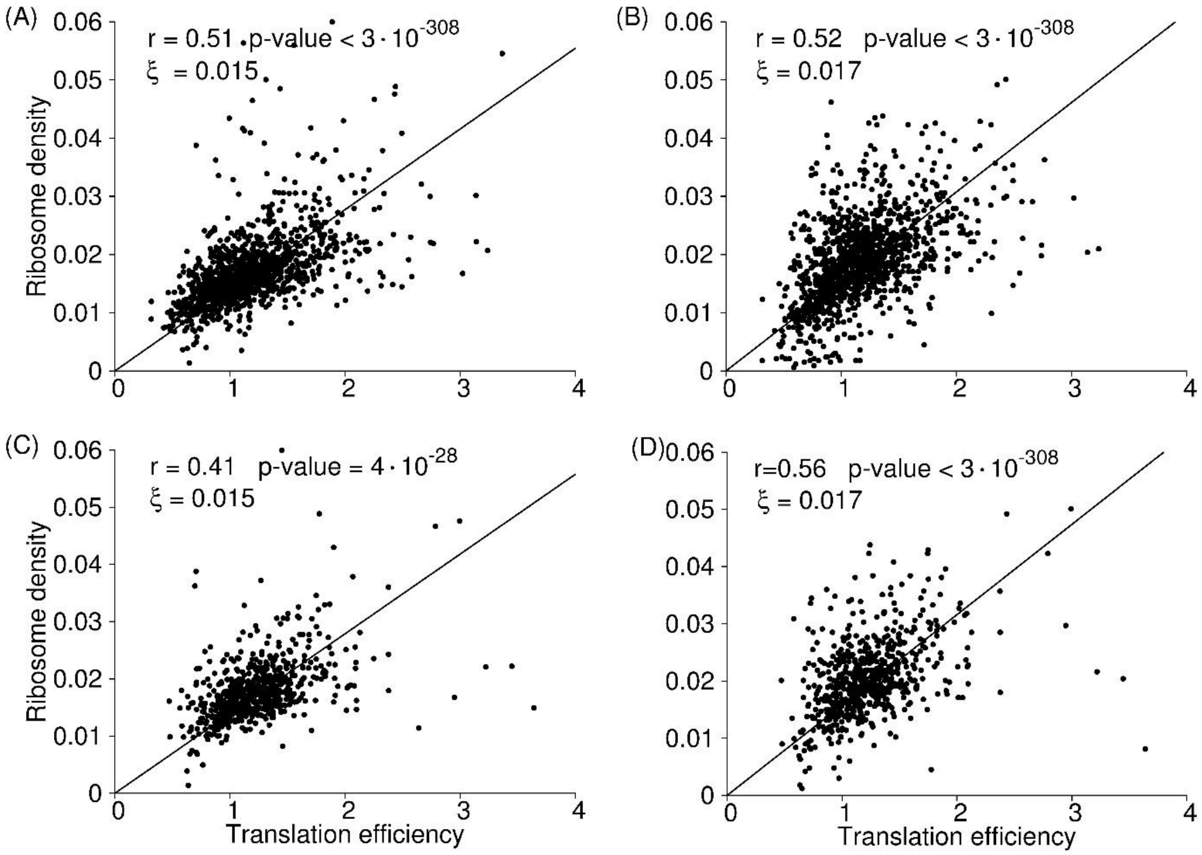


**Figure S7**: **Average ribosome density on a transcript as a function of translation efficiency**. Translation efficiency in (**A**) and (**B**) are calculated using the ribosome profiling and RNA-Seq data reported in Ref. [3]; Translation efficiency in (**C**) and (**D**) are calculated using ribosome profiling and RNA-Seq data reported in Ref. [6]. Ribosome density used in (**A**) and (**C**) are from the polysome profiling data reported in Ref. [5] whereas the ribosome density in (**B**) and (**D**) are provided by Ref. [7]. The solid line in all these figures represent the best fit of $y = \xi x$ line.

| Fragment Size | Frame 0 | Frame 1 | Frame 2 |
|---|---|---|---|
| **24** | **15** | 15/12 | 18/12 |
| **25** | **15** | 12/15 | **18** |
| **26** | 15/12 | 18/15 | 18/15 |
| **27** | **15** | **15** | **18** |
| **28** | **15** | **15** | **18** |
| **29** | **15** | 15/18 | **18** |
| **30** | **15** | **18** | **18** |
| **31** | **15** | **18** | **18** |
| **32** | 18/15 | **18** | **18** |
| **33** | **18** | **18** | **18** |
| **34** | **18** | **18** | 18/21 |

**Table S1: Nucleotide offsets for A-site location from the 5′ end as a function of fragment size and frame.** This offset table was obtained by applying a Linear Programming algorithm to very high coverage pooled dataset of several *S. cerevisiae* ribosome profiling experiment data. In this algorithm, for a fragment size and codon frame, we determine the offset which maximizes the number of reads between the second and the stop codon as ribosome's A-site can only occupy these positions. For certain fragment size and frames, there is ambiguity in identifying the most probable offset therefore, we list the top two offset values here. Reads from these combinations are ignored for calculation of A-site locations.

| Dataset Refs | Total transcripts with initiation rates | Number of transcripts (uAUG) | Number of transcripts (non uAUG) | Transcripts with no 5' UTR annotation | Median initiation rate (uAUG) ($s^{-1}$) | Median initiation rate (non uAUG) ($s^{-1}$) | Mann Whitney U test p-value |
|---|---|---|---|---|---|---|---|
| [3] and [5] | 1287 | 45 | 997 | 245 | 0.095 | 0.112 | **0.006** |
| [3] and [7] | 1249 | 42 | 972 | 235 | 0.115 | 0.134 | **0.009** |
| [6] and [5] | 652 | 16 | 529 | 107 | 0.104 | 0.119 | 0.052 |
| [6] and [7] | 643 | 17 | 518 | 108 | 0.121 | 0.145 | **0.025** |

**Table S2: Transcripts containing at least one upstream AUG (uAUG) have lower median translation initiation rates.** For all possible combinations of ribosome profiling and RNA-Seq data [3,6] and polysome profiling data [5,7], we see a consistent result that the median translation initiation rate is lower for transcripts with at least one uAUG. The result is not statistically significant for combination of Refs. [6] and [5] (p-value =0.052)

| Dataset Refs | Number of transcripts (similar to Kozak sequence) | Number of transcripts (dissimilar to Kozak sequence) | Median initiation rate (similar to Kozak sequence) ($s^{-1}$) | Median initiation rate (dissimilar to Kozak sequence) ($s^{-1}$) | Mann Whitney U test p-value |
|---|---|---|---|---|---|
| [3] and [5] | 201 | 202 | 0.116 | 0.100 | **0.005** |
| [3] and [7] | 197 | 193 | 0.138 | 0.123 | **0.012** |
| [6] and [5] | 127 | 74 | 0.121 | 0.112 | **0.034** |
| [6] and [7] | 126 | 70 | 0.146 | 0.136 | 0.065 |

**Table S3: Transcripts with sequence context around start codon similar to Kozak sequence have higher translation initiation rates.** For all possible combinations of ribosome profiling and RNA-Seq data [3,6] and polysome profiling data [5,7], we see a consistent result that the median translation initiation rate is higher for transcripts with sequence context similar to Kozak sequence (see Methods for details). The result for combination of Refs [6] and [7] is however not statistically significant (p-value = 0.065)
